# *Leishmania amazonensis* controls macrophage-regulated cell death to establish chronic infection *in vitro* and *in vivo*

**DOI:** 10.1101/2022.09.14.507851

**Authors:** Hervé Lecoeur, Sheng Zhang, Hugo Varet, Rachel Legendre, Caroline Proux, Capucine Granjean, Philippe Bousso, Eric Prina, Gerald F. Späth

## Abstract

Pathogenic protists of the genus *Leishmania* have evolved various strategies to exploit macrophages as host cells and subvert their immuno-metabolic functions to favour intracellular parasite survival. Surprisingly little is known on how *Leishmania* affects regulated cell death (RCD) pathways of its host cell, even though increased survival of *in vitro* infected macrophages has been reported, and chronic macrophage infection *in vivo* causes the devastating immunopathologies of leishmaniasis. To overcome this limitation and gain first systems-level insight into the interaction between intracellular *Leishmania* and the host cell RCD pathways, including apoptosis, pyroptosis and necroptosis, we applied transcriptomic analyses on *L. amazonensis*-infected, primary macrophages (termed LIMs) and used YO-PRO-1 to monitor cell death by fluorescent microscopy. RNAseq analyses at day 3 post-infection (PI) revealed dichotomic dysregulation of more than 60% of RCD-related genes in LIMs, characterized by up-regulation of anti-RCD and down-regulation of pro-RCD markers, including key regulators common to the three forms of cell death such as *casp8, fadd, tradd, tnfaip3, tax1bp1, birc3*, and *itch*. This profile correlated with expression changes of transcription factors known to regulate RCD, including AP1 and NF-κB family members, *pparγ* and *cebpβ*. Consequently, LIMs showed remarkable longevity in culture for at least 50 days, despite a constant increase of parasite burden to about 100 parasites per cell, while non-infected cells were cleared from the culture in just a few days. Longitudinal expression analysis of LIMs at days 0, 3, 15, and 30 PI by RT-qPCR confirmed stable maintenance of this high longevity profile with the dichotomic decrease and increase of RCD-activators and -inhibitors, respectively. LIMs further showed significant resistance to RCD-inducing signals compared to non-infected cells, including CSF-1 deprivation (intrinsic apoptosis), actinomycin D treatment (extrinsic apoptosis), LPS/ATP stimulation (pyroptosis). Significantly, we extended the anti-RCD expression pattern and RCD resistance phenotype to *L. amazonensis*-infected macrophages recovered from lesions, thus validating our long-term *in vitro* infection system as an easily accessible model to study chronic macrophage infection. In conclusion, our analyses firmly document the pan-anti RCD effect of *L. amazonensis* on its macrophage host cell *in vitro* and *in vivo* and shed important new light on mechanisms underlying *Leishmania* chronic infection.

## INTRODUCTION

One key strategy shared between intracellular pathogens is their capacity to prolong host cell survival by hijacking pathways controlling cell death. This strategy not only safeguards the pathogens’ metabolic niche, but also avoids exposure to anti-microbial activities triggered by dying cells. This phenomenon is very well illustrated by intracellular parasites of the genus *Leishmania* that infect innate immune system such as macrophages (Mϕs) and dendritic cells (Liu and Uzonna, 2012) and exploits the phenotypic plasticity of these cells to establish a unique immuno-metabolic phenotype that favour parasite survival and chronic infection (Arango Duque and Descoteaux, 2015; Ferreira et al., 2021; Lecoeur et al., 2021; Saunders and McConville, 2020; Van Assche et al., 2011; Zhang et al., 2022).

In response to intracellular microbial infections, Mϕs deploy an arsenal of potent anti-microbial activities (nitric oxide, reactive oxygen intermediates, anti-microbial peptides) and restrict the microbes’ proliferation by limiting its access to essential nutrients (Weiss and Schaible, 2015). Host cell suicide represents the most extreme outcome of the anti-microbial response, which can trigger immune recognition and elimination of the exposed pathogen (Behar et al., 2011; Chow et al., 2016; Robinson et al., 2019). The process of self-destruction is tightly controlled through well-defined pathways of Regulated Cell Death (RCD) (Galluzzi et al., 2018; Jorgensen et al.; Riera Romo, 2021), including apoptosis, pyroptosis and necroptosis, all of which can be associated with microbial infections (Hacker, 2018; Tummers and Green, 2022; Xia et al., 2020). Each of these RCD pathways is characterized by specific activation signals that trigger defined signalling cascades, and their link to either the pro- or anti-inflammatory host response (Riera Romo, 2021; Tang et al., 2019). Apoptosis can be triggered by growth factor deprivation (intrinsic pathway) and by death factors of the TNF family (such as FasL, APO-2L/-3L) or actinomycin D (extrinsic pathway) (Galluzzi et al., 2018; Liu et al., 2016; Merkel et al., 2012). This form of RCD is orchestrated by members of the caspase family that carry cysteine protease activity inducing specific proteolysis and cell destruction. Apoptosis is the typical non-lytic type of RCD that avoids and even suppresses inflammatory responses by i) releasing anti-inflammatory metabolites to modulate gene expression in neighbouring cells (Medina et al., 2020), ii) secreting “come-get-me” molecules (Grimsley and Ravichandran, 2003), and iii) expressing “eat-me” signals (Grimsley and Ravichandran, 2003) that trigger their swift removal by macrophages (efferocytosis) before releasing their intracellular content(Grimsley and Ravichandran, 2003). *In vivo*, engulfment of apoptotic cells modulates the phagocyte’s immune-metabolism to resolve inflammation (Birge and Ucker, 2008; Kourtzelis et al., 2020).

In contrast, necroptosis and pyroptosis are two regulated necrotic (i.e. lytic) processes that have been associated to potent inflammatory responses *in vivo* (Bergsbaken et al., 2009; Frank and Vince, 2019; Grootjans et al., 2017; Pasparakis and Vandenabeele, 2015). Both forms of RCD trigger the formation of pores in the plasma membrane (Kolbrink et al., 2020) that allows the release of Damage-Associated Molecular Patterns (DAMPs) (Frank and Vince, 2019; Murao et al., 2021) and in the case of pyroptosis, the release of potent pro-inflammatory cytokines such as IL1β (Heilig et al., 2018). Pyroptosis is mediated by inflammasome activation in response to pathogenic agents or various stimuli such as toxins or ATP via intracellular receptors such as NOD-like receptor family, pyrin domain containing 3 (NLRP3) and absent in melanoma 2 (AIM2), which cause gasdermin-dependent cell lysis (Paik et al., 2021; Yu et al., 2021). Necroptosis is a form of regulated necrosis dependent on receptor-interacting serine/threonine kinase 1 and/or 3 (RIPK1, RIPK3) and mixed lineage kinase domain-like pseudokinase (MLKL) (Grootjans et al., 2017). It has been initially identified as a backup death pathway in response to inhibition of caspase-8-dependent apoptosis (Holler et al., 2000), but can also be induced by different mediators or pathogens (Robinson et al., 2019). Despite their seemingly different characteristics, apoptosis, pyroptosis and necroptosis are interconnected and co-regulated via key shared regulatory molecules (Bedoui et al., 2020; Place et al., 2021), a redundancy that likely evolved to overcome pathogen-induced suppression of individual RCD pathways (Tummers and Green, 2022).

Several studies showed that *Leishmania* spp can dampen individual RCD pathways in infected macrophages (Akarid et al., 2004; Donovan et al., 2009; Lisi et al., 2005; Moore et al., 1994; Rodriguez-Gonzalez et al., 2016). Several mechanisms were identified, such as increased expression of Cellular Inhibitors of Apoptosis Proteins (cIAPs), differential expression of B-cell lymphoma 2 (BCL2) family members, decreased expression of key RCD effectors (pro-caspase-1, RIKP3, PGAM5 and MLKL) or changes in PI3K/AKT signalling (Cianciulli et al., 2018; Dashzeveg and Yoshida, 2015; Giri et al., 2016; Luz et al., 2018; Pandey et al., 2016; Roy et al., 2017; Ruhland et al., 2007; Saha et al., 2019a; Saha et al., 2019b). However, our current knowledge on *Leishmania*-mediated RCD subversion is limited to only short-term infections, the exclusive use of *in vitro* infection systems, and the lack of integrative analyses that allow mechanistic insight into all major RCD pathways during infection. Here we overcome these limitations and gain first systems-level insight into the interaction between intracellular *Leishmania amazonensis* (*L. am*) and apoptotic, pyroptotic and necroptotic host cell pathways. We provide first evidence that *Leishmania*-infected macrophages (termed LIMs) show a remarkable longevity in culture of over 50 days, which was linked to profound expression changes in transcription factors known to regulated RCD and increased resistance to the various forms of cell death. Significantly, both expression changes and cell death-resistant phenotype were validated in *in situ*-infected macrophages directly isolated from mouse footpad lesions. Our data confirm our previous published observations on *Leishmania*-dependent changes in macrophage nuclear activities and cell survival (Lecoeur et al., 2021; Lecoeur et al., 2020a; Lecoeur et al., 2020b), and propose our long-term *in vitro* infection system as a useful correlate for chronic infection to investigate parasite and host factors involved in cell death regulation.

## MATERIAL AND METHODS

### Ethics statement

Six-week-old female C57BL/6 and Swiss *nu/nu* mice were purchased from Janvier (Saint Germain-sur-l’Arbresle, France). Rag2^tm1Fwa^ (Shinkai et al., 1992) were bred at Institut Pasteur. All animals were housed in A3 animal facilities according to the guidelines of Institut Pasteur and the “Comité d’Ethique pour l’Expérimentation Animale” (CEEA) and protocols were approved by the “Ministère de l’Enseignement Supérieur; Direction Générale pour la Recherche et l’Innovation” under number #19683.

### Isolation and culture of bone marrow-derived macrophages

Bone marrow cell suspensions were recovered from tibias and femurs of C57BL/6 mice in DMEM medium (Gibco, Life technologies) and cultured in medium complemented with mouse recombinant colony-stimulating factor 1 (mrCSF-1, ImmunoTools) (Lecoeur et al., 2020a). Precursors were seeded for one night at 37°C in a 7.5% CO2 atmosphere at 3 × 10^7^ cells / 12 ml of complete medium containing 50 ng/ml of rm-CSF-1, in Petri dishes for tissue culture (Falcon® 100 mm TC-treated Cell Culture Dish ref OPTILUX 353003) for the removal of unwanted adherent cells. Unattached cells were recovered and cultured at one million cells per ml in non-treated Petri dishes (Greiner bio-one 664161) at 37°C in a 7.5% CO2 atmosphere for 6 days with 50 ng/mL mrCSF-1. After detachment with 25 mM EDTA, 2 million Bone Marrow-Derived Macrophages (BMDMs) were seeded per well in 12-well plates in 2 ml culture medium containing 30 ng/mL mrCSF-1 (for fluorescence microscopy and western blot analyses) or in 10 ml in petri dishes (for RNA isolation and RNAseq analyses). Analyses were performed from day 1 to day 48 post-infection. For long-term experiments, half of the medium was replaced by complete medium supplemented with 30 ng/mL mrCSF-1 once a week.

### Parasite isolation and generation of *in vitro Leishmania-*infected macrophages (*iv*LIMs)

mCherry transgenic, tissue-derived amastigotes of *Leishmania amazonensis* strain LV79 (WHO reference number MPRO/BR/72/M1841) were isolated from infected footpads of Swiss nude mice (Lecoeur et al., 2010). Amastigotes were added at a ratio of 4 amastigotes per macrophage, yielding an infection efficiency of 95% at 3 days post-infection (PI) as judged by fluorescence microscopy analysis of Hoechst-stained samples. LIMs were cultured at 34°C for 3 to 50 days PI. Long-term cultures were maintained by replacing the supernatant once a week with a fresh complete medium containing 30 ng/ml mrCSF-1.

### Isolation of *ex vivo Leishmania-*infected macrophages (*ev*LIMs) from mouse lesions

*ev*LIMs were isolated from footpad lesions of Rag2^tm1Fwa^ mice obtained 2 months after the inoculation of amastigotes. Mice were euthanized by carbon dioxide inhalation, footpads were removed and placed on a 100 μm cell strainer in treated with Digestion Buffer (DB) containing 50 U/ml DNAse I (Sigma-Aldrich), 100 U/ml collagenase II (Sigma-Aldrich), 100 U/ml collagenase IV (Sigma-Aldrich) and 1 U/ml dispase II (Roche Applied Science) in DMEM. One ml of DB was perfused in the footpad at 5 different locations for 30 minutes at 34°C. This step was repeated two times. *ev*LIMs were recovered and washed (centrifugation at 50 *g*, 10 min, 4 °C) before processing. For *ev*LIMs culture, cells were placed for 1 hour at 34°C in Petri dishes (Falcon® 100 mm TC-treated Cell Culture Dish ref OPTILUX 353003) to remove free amastigotes potentially released from cells damaged during the procedure. The supernatant containing *ev*LIMs was then used for further culture.

### RCD induction in *in vitro* and *ex vivo* macrophages

Apoptosis was induced by rm-CSF1 deprivation (from 30 ng/ml to total removal) for different periods of time or by the addition of 5μg/ml Actinomycin D (AD, #A1410, Sigma-Aldrich) for 16 hours (Lisi et al., 2005). Pyroptosis was induced after inflammasome priming and activation by sequential stimulation with 500 ng/ml lipopolysaccharides from *Escherichia coli* O111:B4 (# LPS11-1, Alpha Diagnostic) for 16 h and 5 mM adenosine triphosphate (ATP, # A26209, Sigma-Aldrich) for the time required for the analysis (1 to 8 hours). For RNAseq analyses, ATP was added for 30 minutes to LPS-treated cells prior to RNA isolation. Experiments were conducted in 12 well plates (0.5 × 10^6^ cells per well). For long-term analyses, 50% of the medium was discarded and replaced by a fresh complete medium supplemented by 30 ng/ml mrCSF1. Apoptosis, pyroptosis and necroptosis were analysed according to recently published guidelines (Hu et al., 2021).

### Epifluorescence microscopy analyses of RCD and automated image analysis

Nuclei from both live and dead macrophages were stained by 1 μM Hoechst 33342 (ThermoFisher Scientific). Nuclei from apoptotic, pyroptotic and necroptotic macrophages were revealed by staining with by 1 μM YO-PRO-1 (ThermoFisher Scientific) (Darzynkiewicz et al., 2011; Idziorek et al., 1995; Rosazza et al., 2020). Both markers were added to the culture medium and cells were incubated for 15 min at room temperature. Image acquisition was done using the EVOS M5000 microscope (ThermoFisher Scientific) for transmitted light, and florescence signals detected at red (mCherry expressing parasites), blue (Hoechst) and green (YO-PRO-1) wavelengths. Five fields were acquired at the 20x objective.

Automatic Image analysis was performed using the ImageJ software package on at least 800 cells. Automated counting of macrophage nucleus was performed by detecting the Hoechst 33342 signal. For morphometric analysis, macrophage area and mCherry intensity information (related to parasite load) were simultaneously determined for each individual macrophage. Graphical representations were generated with the GraphPad Prism software package.

### RNA extraction and transcriptional analyses by RNAseq and reverse transcription quantitative PCR analyses

RNA isolation was performed with the RNeasy^+^ isolation kit (Qiagen) according to the manufacturer’s instructions. Evaluation of RNA quality was carried out by optical density measurement using a Nanodrop device (Kisker, http://www.kisker-biotech.com) as previously described (Lecoeur et al., 2020a). Total RNA was isolated from 3 independent biological replicates of uninfected and *Leishmania*-infected macrophages.

DNAse-treated RNA extracts were processed for library preparation using the Truseq Stranded mRNA sample preparation kit (Illumina, San Diego, California) according to the manufacturer’s instructions. An initial poly(A)^+^ RNA isolation step (included in the Illumina protocol) was performed with 1 μg of total RNA to isolate the mRNA fraction and remove ribosomal RNA. The mRNA-enriched fraction was fragmented by divalent ions at high temperatures. The fragmented samples were randomly primed for reverse transcription followed by second-strand synthesis to create double-stranded cDNA fragments. No end repair step was necessary. An adenine was added to the 3’-end and specific Illumina adapters were ligated. Ligation products were submitted to PCR amplification. The quality of the obtained libraries was controlled using a Bioanalyzer DNA1000 Chips (Agilent, # 5067-1504) and quantified by spectrofluorimetric analysis (Quant-iT™ High-Sensitivity DNA Assay Kit, #Q33120, Invitrogen).

Sequencing was performed on the Illumina Hiseq2500 platform to generate single-end 65 bp reads bearing strand specificity. Reads were cleaned using cutadapt version 1.11 and only sequences at least 25 nt in length were considered for further analysis. STAR version 2.5.0a (Dobin et al., 2013), with default parameters, was used for alignment on the reference genome (GRCm38 from Ensembl database 94). Genes were counted using featureCounts version 1.4.6-p3 (Liao et al., 2014) from the Subreads package (parameters: -t gene -s 0). Transcriptomic data are made publicly available at the NCBI’s Gene Expression Omnibus repository (Superseries GSE205860). The RCD-related gene list is provided for each pathway in Supplementary figure 1.

RT-qPCR was carried out in 384-well PCR plates (Framestar 480/384, 4titude, Dominique Dutscher) using the iTaq Universal SYBR® Green Supermix (Bio-Rad) and 0.5 μM primers with a LightCycler® 480 system (Roche Diagnostics, Meylan, France). Primer information for all targets is detailed in Supplementary table S1. Crossing Point values were determined using the second derivative maximum method (LightCycler® 480 Basic Software). The relative expression software tool (REST©-MCS) (Pfaffl et al., 2002) was used to calculate fold change (FC) values. The list of primers for each analyzed target gene is provided in Supplementary table 2. Normalization was performed with the geometric mean of the two best reference genes *ppih* and *mau2* as determined by the GeNorm and Normfinder programs (data not shown). For statistical analysis of gene expression levels, Cp values were first transformed into relative quantities (RQ) and normalized (Lecoeur et al., 2020a). Nonparametric Kruskal-Wallis tests were performed on Log transformed Normalized Relative Quantity values. Results were expressed as Z-scores calculated by relative expression compared to the day 0 unpolarized group (long-term analyses). Heatmaps of Z-scores were generated (R, version 4.1.2) using homemade scripts based on native graphical functions.

### *In silico* analyses

String network analyses were performed using Cytoscape V3.9.1, the interaction of TFs and targets were extracted from Transcriptional regulator relationships unraveled by a sentence-based text mining database (TRRUST v2: an expanded reference database of human and mouse transcriptional regulatory interactions (Han et al., 2018). TF-targets gene networks were obtained with the ENCODE transcription Factor targets dataset.

### Statistical analysis

Statistical analysis of gene expression levels determined by RT-qPCR was performed using SigmaPlot Software (SigmaPlot for Windows Version 11.0, Build 11.2.0.5). Other statistical analyses were performed by the nonparametric Wilcoxon rank-sum test using the GraphPad Prism 7.03 software.

## RESULTS AND DISCUSSION

### *Leishmania amazonensis* infected macrophages show an important increase in longevity

Previous studies correlated short-term *Leishmania* infection with protection from cell death (Akarid et al., 2004; Donovan et al., 2009; Lisi et al., 2005; Moore et al., 1994; Rodriguez-Gonzalez et al., 2016). Here we assessed the ability of primary macrophages to survive chronic intracellular infection with *L. amazonensis* (*L. am*), which to our knowledge was not investigated before. C57BL/6 BMDMs were infected with lesion-derived *L. am* amastigotes expressing mCherry (referred to as ‘*in vitro Leishmania*-infected macrophages’, short ‘*iv*LIMs’). In contrast to non-infected macrophages, *iv*LIMs were able to survive in culture for a long time (more than 30 days) despite a substantial increase in intracellular parasite burden that attained over 70 amastigotes per macrophage after only two weeks of infection as judged by increased mCherry fluorescence intensity and the number of Hoechst 33342-stained parasite nuclei (Figure 1A1 and Supplementary Figure 1). This surge in parasite load correlated with the generation of large communal parasitophorous vacuoles and a substantial increase in mean macrophage area from 1789 to 14679 sq. pixels (Figure 1A2 and 3). Significantly, parasite burden and cell swelling of *iv*LIMs in long-term cultures closely matched infected LIMs purified from the tissue of infected RAG2-KO mice and analyzed *ex vivo* (termed *ev*LIMs) (Figure 1B). Given the very low replication rate of intracellular amastigotes shown *in vivo* (Kloehn et al., 2015), these data also strongly support the increased lifespan of *ev*LIMs.

**Figure 1:**
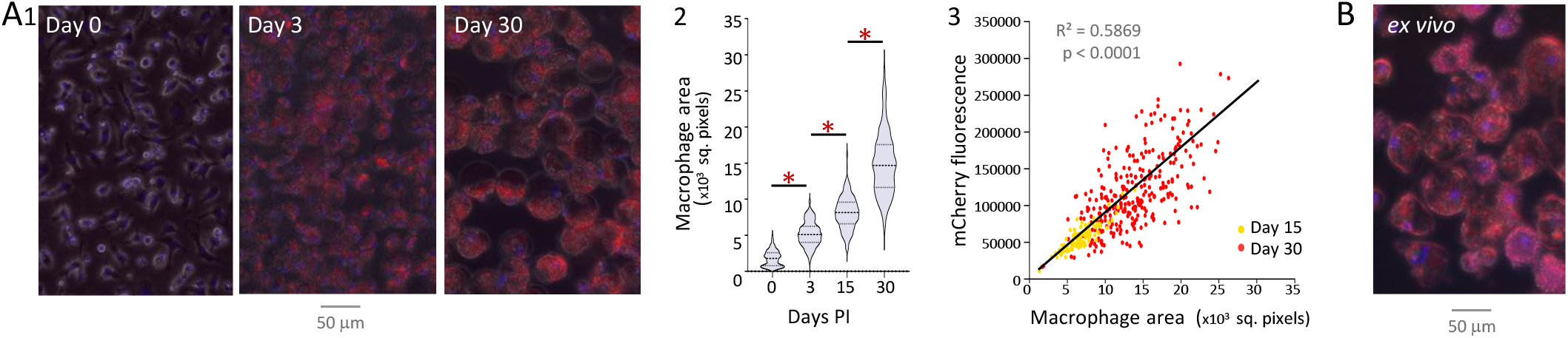
BMDMs *in vitro* and macrophages *in vivo* are permissive to chronic infection by *L. amazonensis* amastigotes. **(A1)** Uninfected BMDMs (day 0) and cells infected with mCherry-transgenic *L. amazonensis* amastigotes at day 3 and 30 post-infection were analyzed by fluorescence microscopy using an EVOS M5000 microscope (20x objective). Macrophage nuclei and parasites were respectively revealed by Hoechst 33342 staining (blue fluorescence) and mCherry expression (red fluorescence). Superpositions of fluorescence and bright field images are shown. **(A2)** Violin plots representing the median and quartiles of BMDM area (squared pixels) during long-term infection. *, p < 0.0001. **(A3)** Correlation between cell area and mCherry fluorescence (raw integrated density) in individual BMDMs at days 15 and 30 post-infection. Linear regression curve, R^2^ and p-value are indicated. **(B)** Lesional macrophages obtained from RAG2-KO mouse 60 days after infection and analyzed *ex vivo* by fluorescence microscopy by Hoechst 33342 staining (blue fluorescence, macrophage nuclei) and mCherry transgenic expression (red fluorescence, parasites).

Together these data demonstrate that long-term *Leishmania*-infected macrophages *in vitro* are a very good correlate to chronically infected macrophages *in vivo*. We assessed in the following if increased longevity is caused by increased resistance of LIMs to RCD processes.

### *L. am* prevents RCD in infected *in vitro* and *in vivo* macrophages

Induction of intrinsic apoptosis by long-term deprivation for the anti-apoptotic growth factor rm-CSF1 (Becker et al., 1987) caused massive cell death in non-infected macrophages as judged by the increased number of YO-PRO-1 positive, apoptotic cells (Figure 2A1a) and a 50% reduction in cell density after three days of treatment (Figure 2A2 and Supplementary Figure 2B, time point “a”). In contrast, less than 5% of *iv*LIMs were stained with YO-PRO-1 at the same time of rm-CSF1 deprivation, with the few apoptotic cells being devoid of intracellular parasites, thus further sustaining the parasite’s anti-apoptotic action, which likely occurs in *cis* and cannot be transferred in *trans* to non-infected bystander cells (Figure 2A1b and 2A2). *iv*LIMs maintained a low apoptosis rate all along the 50-day culture (Figure 2A1b, c, and Figure 2A2) and displayed a 50% drop in cell density only after 26 days PI (Supplementary Figure 2B, time point “b”). This remarkable survival phenotype was confirmed in *ev*LIMs that persisted *ex vivo* for 7 days in the absence of rm-CSF1 (Figure 2Ad and Supplementary Figure 2C) despite their high parasite load and the physical stress associated with their purification from infected tissue. A similar phenotype in *iv*LIMs and *ev*LIMs was observed when triggering extrinsic apoptosis by 16-hour exposure to actinomycin D (AD) stimulation (Figure 2A3). Likewise, induction of NLRP3-dependent pyroptosis by LPS/ATP stimulation (Lecoeur et al., 2020a; Yu et al., 2021) caused substantial cell death in non-infected control cells (Figure 2B1a), attaining 80% of YO-PRO-1 positive cells at 8 hours of ATP stimulation (Figure 2B2), while only 40% of dead cells were observed in *iv*LIMs (Figure 2B1b, c and Figure 2B2) and in *ev*LIMs (Figure 2B3).

**Figure 2:**
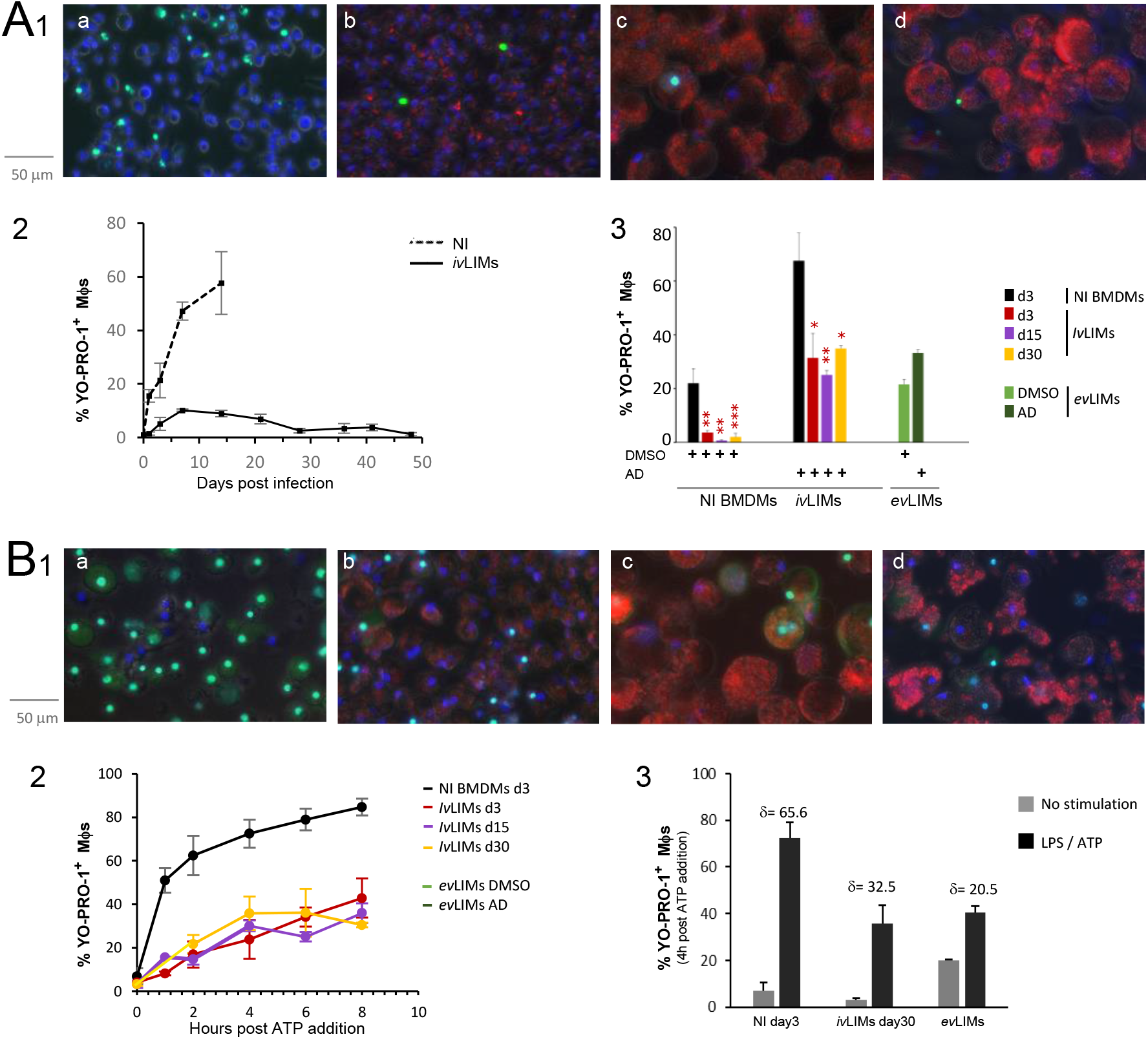
*In vitro* and *in vivo* chronically infected macrophages show a global resistance to RCD processes. **(A1)** Cells were exposed to rm-CSF1 deprivation to induce intrinsic apoptosis. Cell death was monitored by YO-PRO-1 staining (green color) and fluorescence microscopy *in vitro* in non-infected (A1a) and *L. amazonensis-*infected BMDMs (*iv*LIMs) at days 3 (A1b) and 30 (A1c) post-infection as well as *ex vivo* in lesional macrophages (*ev*LIMs) 7 days after isolation (A1d, see legend of Figure 1 for details). Analyses were performed using an EVOS M5000 microscope with a 20x objective. Macrophage nuclei were stained with Hoechst 33342 (blue color) and parasites were revealed by mCherry (red color). **(A2)** Percent of YO-PRO-1 positive, apoptotic cells during long-term rm-CSF1 deprivation in non-infected (NI, dotted line) and infected (*iv*LIMs, continuous line) macrophages is shown. Macrophage numbers were estimated microscopically based on the Hoechst 33342 signal (n = 5 fields). **(A3)** Percent of YO-PRO-1 positive cells in presence of DMSO (vehicle control) and following induction of extrinsic apoptosis by 16-hour treatment with Actinomycin D (AD)of *iv*LIMs at days 3, 15 and 30 PI and of *ev*LIMs. Mean values and SEM for three independent experiments are shown. Statistical differences between non-infected and infected groups are indicated by the corresponding p values: *, p < 0.03; **, p < 0.005; ***, p < 0.0001. **(B1)** Pyroptosis was triggered in BMDMs by overnight LPS stimulation and 4h ATP treatment, and cell death was monitored by YO-PRO-1 staining in non-infected controls (NI, B1a), *iv*LIMs after 3 days (B1b) and 30 days of infection (B1c) and in *ev*LIMs right after their isolation from infected footpads (dB1). **(B2)** Eight-hour kinetic analysis of pyroptosis in *iv*LIMs induced at days 3, 15 and 30 post-infection. **(B3)** Pyroptosis rates in non-infected BMDMs (NI), *iv*LIMs and *ev*LIMs in response to LPS stimulation and 4-hour-ATP treatment. The difference in YO-PRO-1 positive cells between infected and non-infected cells is indicated by the δ values.

In conclusion, we provide here first evidence for active, *Leishmania*-driven pan-inhibition of macrophage RCD pathways, reminiscent of other intracellular pathogens such as cytomegalovirus that inhibit apoptosis and necroptosis (Brune and Andoniou, 2017) or *Coxiella burnetii* that interferes with host cell apoptosis and pyroptosis (Cordsmeier et al., 2019).

### The anti-RCD phenotype of *iv*LIMs is controlled at the transcriptional level

To gain first insight into parasite subversion strategies that can be linked to host cell longevity, we applied RNAseq analysis on uninfected BMDMs and *iv*LIMs at day 3 PI and monitored the effect of infection on genes involved in different RCD pathways. As judged by the Odds ratios and False Discovery Rates (FDR), more than 60% of the genes linked to apoptosis, pyroptosis and necroptosis show significant expression modulation (Figure 3A1). Mapping these changes across different categories of genes known to promote or inhibit RCD revealed a regulatory dichotomy that causes dual inhibition via (i) down-modulation of pro-RCD genes that encode for surface receptors, signaling molecules, and components of key RCD complexes, and (ii) up-regulation of caspases, inflammasome and necrosome inhibitors (Figure 3A2). We further revealed differential expression in LIMs of key regulators that are shared across all RCD pathways, including down-regulation of *caspase-8* and *fadd*, and up-regulation of the two potent RCD inhibitors *tax1bp1* and *tnfaip3* (Figure 3A3), the latter one known to counteract apoptosis, pyroptosis and necroptosis (Priem et al., 2020). This expression pattern was confirmed by RT-qPCR in both *iv*LIMs from long-term infection and *ev*LIMs obtained from infected lesions, the latter one showing the strongest expression of *tax1bp1* and *tnfaip3* (Figure 3B).

**Figure 3:**
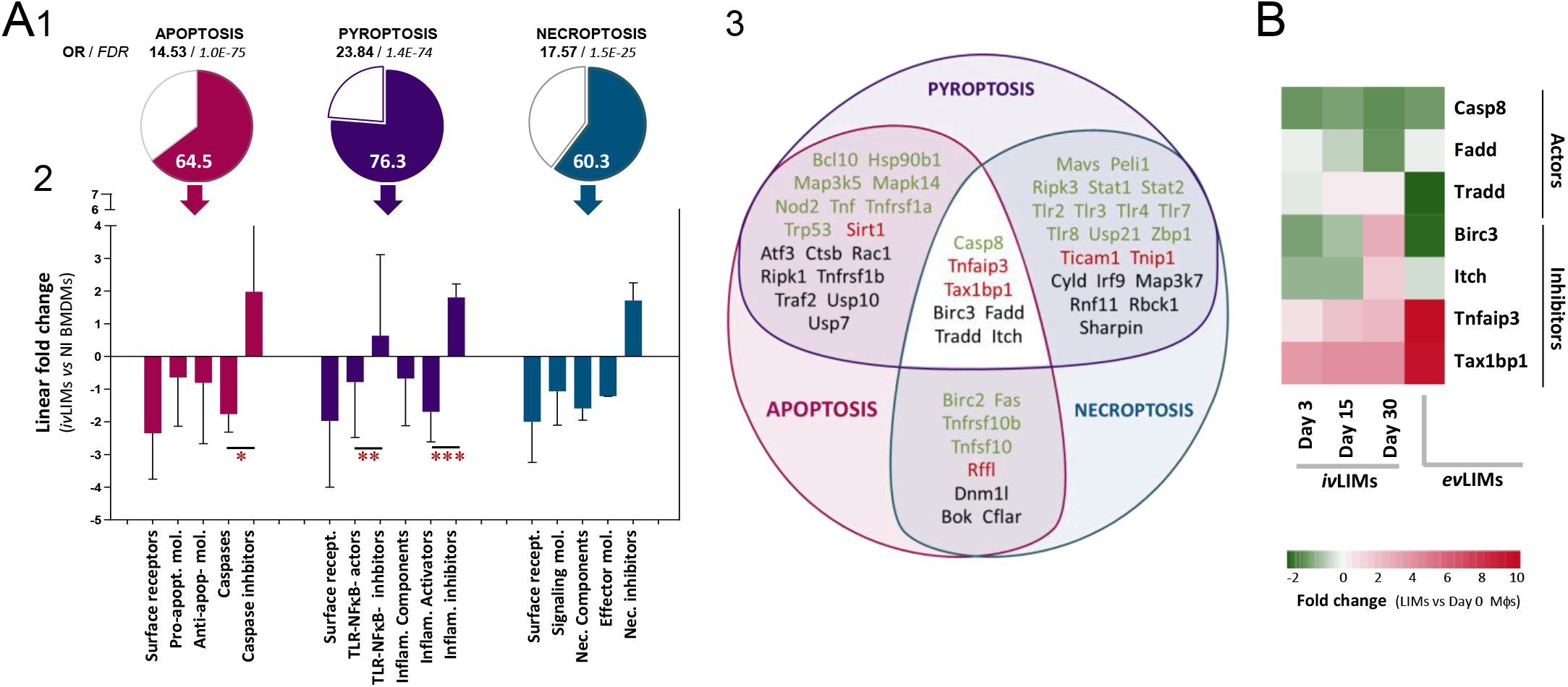
Transcriptomic inhibition of RCD pathways in *iv*LIMs and *ev*LIMs. **(A)** RNAseq analysis of RCD genes (Supplementary table 1) in non-infected BMDMs and *iv*LIMs at day 3 post-infection. The percentage of regulated genes implicated in apoptosis, pyroptosis and necroptosis (A1) and the fold expression change for different categories of genes implicated in RCD (A2) are shown. RNAseq-based global transcriptomic analysis related to apoptosis, pyroptosis (and also necroptosis). Odds Ratios (OR) and False Discovery Rates (FDR) values for each pathway are indicated. *, p < 0.051; **, p < 0.04; ***, p < 0.02. The Venn diagram (A3) indicates expression changes in *iv*LIMs compared to non-infected BMDMs for the shown genes that are either specific or shared between the different RCD pathways. The letter color corresponds to the observed expression changes (p < 0.05): red, increased abundance; green, reduced abundance; black, no change. **(B)** Heat map of RT-qPCR analysis of the common RCD regulators (see center of the Venn diagram shown in A3) in *iv*LIMs at days 3, 15 and 30 post-infection, and lesion-derived *ev*LIMs. Expression changes were calculated relative to non-infected control BMDMs.

In conclusion, our data uncover a regulatory dichotomy that inhibits host cell RCD by inhibiting pro- and at the same time promoting anti-RCD gene expression. Increased expression of *tnfaip3* was previously observed in *L. am* infected BALB/c BMDMs (Lecoeur et al., 2020a) and *L. infantum* infected THP1, RAW 264.7 cells and BMDMs (Gatto et al., 2020; Roy et al., 2017), suggesting this factor as a key regulator in pan-RCD inhibition during *Leishmania* infection. We next investigated the effect of *L. am* infection on the different RCD pathways by mapping our RNAseq results onto the apoptotic, pyroptotic and necroptotic pathways, and validating observed expression changes in more detail by RT-qPCR analyses of *iv*LIMs from long-term infection and *ev*LIMs obtained from infected lesions.

### Dichotomic inhibition of apoptotic pathways in *iv*LIMs and *ev*LIMs

Mapping our expression results on the apoptotic pathways confirmed the dichotomic regulation we observed during *L. am* infection (Figure 4A1 and A2). Several expression changes explain the observed inhibition of extrinsic apoptosis, including down-modulation of (i) the surface receptors *fas, tnfrsf1a*, and *tnfrsf10b*, (ii) the adaptor molecules *traf5* and *tradd*, and (iii) the effector caspases *casp3* and *casp6*. This pathway is further inhibited by the up-regulation of *tnfaip3/tnip1/taxbp1*, the caspase inhibitor *birc6*, and the inhibitor of caspase-activated DNase *dffa* (Figure 4A1). We further observed decreased expression of several components of the AP1 transcription factor complex (i.e. *fos, fosb, fosl1, junb*), which possibly leads to the decreased expression of pro-apoptotic effectors such as *bcl2l11 (bim)* (Whitfield et al., 2001) (Figure 4A1). A similar regular dichotomy was observed for intrinsic apoptosis showing (i) down-modulation of pro-apoptotic *igf1r* and the caspases *casp12, casp2*, and *casp8*, and (ii) up-regulation of *aven* that stabilizes the anti-apoptotic molecule *bcl2l1* and inhibits Apoptotic Protease Activating Factor 1 (APAF1) known to form one of the central hubs in the apoptosis regulatory network that triggers caspase activation (Chau et al.; Kutuk et al., 2010). Conversely, we further observed expression changes that did not fit this dichotomy, such as (i) down-modulation of the anti-apoptotic *Bcl2* molecule that was previously observed in *L. mexicana-*infected DCs (Rodriguez-Gonzalez et al., 2022), or (ii) up-regulation of pro-apoptotic BID, which however may not be cleaved to its active form since *iv*LIMs show reduced expression of *calpain1* (*capn1)* and components of the PIDDosome - a multiprotein complex that drives activation of caspase-2 (Figure 4A1). Importantly, *L. am* infection also seems to prevent a key step in apoptotic cell death represented by Mitochondrial Outer Membrane Permeabilization (MOMP) (Dadsena et al., 2021), which is a key checkpoint in apoptosis that activates the caspase cascade. The induction of MOMP is regulated by pro-apoptotic members of the BCL2 family that were down regulated in *iv*LIMs (i.e. *bax, pmaip1, bcl2l11, bbc3*), and anti-apoptotic *mcl1* that was upregulated in *iv*LIMs as previously observed in other experimental *Leishmania* infection systems (Giri et al., 2016; Ruhland et al., 2007).

**Figure 4:**
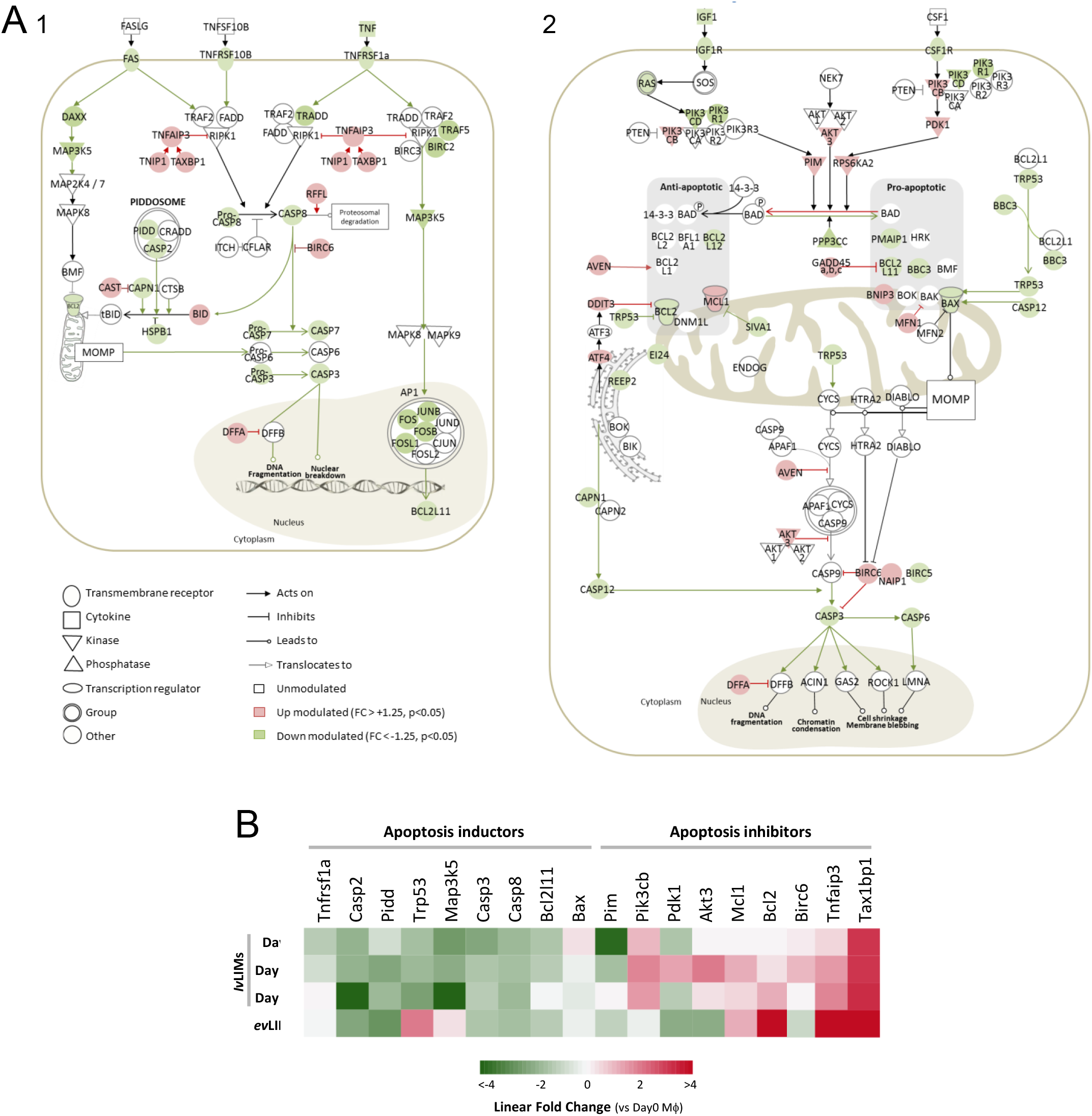
The anti-apoptotic phenotype of *iv*LIMs and *ev*LIMs correlates with dual transcriptomic repression of the corresponding pathway. **(A)** Subversion of extrinsic (A1) and intrinsic (A2) apoptosis pathways in *iv*LIMs at day 3 post-infection as assessed by RNAseq analysis. Significantly increased (linear FC > +1.25) and decreased (linear FC < -1.25) mRNA abundances are depicted in red and green color, respectively. **(B)** Heatmap of transcriptomic changes observed by RT-qPCR analysis in *iv*LIMs at days 3, 15 and 30 PI (n = 3 independent experiments) and in lesion-derived *ev*LIMs (n = 4 RAG2-KO mice). The analysed transcripts were selected based on our RNAseq results obtained with *iv*LIMs on day 3 PI.

We validated these expression changes observed at day 3 PI in *iv*LIMs from long-term infection and *ev*Lims obtained from mouse lesions using RT-qPCR (Figure 4B). Significantly, in contrast to short-term infections (day 3 PI), both *iv*LIMs at day 30 and *ev*LIMs obtained after two months of *in vivo* infection showed increased expression of the anti-apoptotic regulator *bcl2*, which further strengthens the anti-apoptotic phenotype in these heavily infected cells (Figure 4B). These fine-tuned, transcriptomic changes in the LIMs RCD pathways are reminiscent of our previous analysis of the anti-inflammatory LIMs phenotype (Lecoeur et al., 2020a) and quite unique compared to other infectious systems that inhibit host cell RCD by (i) direct degradation of pro-apoptotic effectors (*Chlamydia trachomatis*) (Pirbhai et al., 2006), ii) activation of the pro-survival kinases Akt and Extracellular signal-Regulated Kinases 1 and 2 (*Coxiella burnetti)* (Voth and Heinzen, 2009), or iii) direct inhibition of the initiator caspase-8 (various viruses) (Mocarski et al., 2011).

### Dichotomic inhibition of the pyroptotic pathway in *iv*LIMs and *ev*LIMs

Pyroptosis represents an inflammasome-induced form of cell death that is promoting microbial elimination (Xia et al., 2019). Our RNAseq results reveal transcriptomic inhibition of this pathway by yet another dichotomic regulation (Figure 5A) that affects key steps in pyroptosis induction. First, NLRP3 inflammasome priming and activation steps were inhibited by (i) down-modulation of activating components such as *eif2ak2, p2rx7, mavs, or ptpn22* (Kelley et al., 2019; Paik et al., 2021) and (ii) up-regulation of the de-activating components *fbxl2, nlrc3, or nlrp10)* (Swanson et al., 2019) of this pathway. These results are in accordance with our previous studies of short-term *L. amazonensis-*infected BALB/c primary dendritic cells and BMDMs (Lecoeur et al., 2020a; Lecoeur et al., 2020b). Second, subsequent NF-κB-mediated, pro-inflammatory responses were inhibited by (i) down modulation of various activators, including surface receptors (i.e. the *cd14/lbp/ly96/tlr4* complex, *il1r1, il18r1/il18rap* and *tnfrsf1a*), the adaptor molecule *myd88* and the protein kinase *mapk14*, and (ii) up-regulation of inhibitors of the pathway such as *tnip1, tnfaip3, tax1bp1, nlrc3, sarm, sirt1* and *sirt2*. The dual inhibition of the TLR/NF-κB signaling pathway was further associated with reduced expression of transcription factors that control the gene expression of different inflammasome components (*trp53, zfp36* and *stat1*) (Christgen et al.; Gupta et al., 2001; Haneklaus et al., 2017) explaining the reduced expression of *nlrp3, nlrc4, nlrp1b, mefv, casp1, pycard* and CASPASE-1 substrates (*il18 and il1β)* in *iv*LIMs.

**Figure 5:**
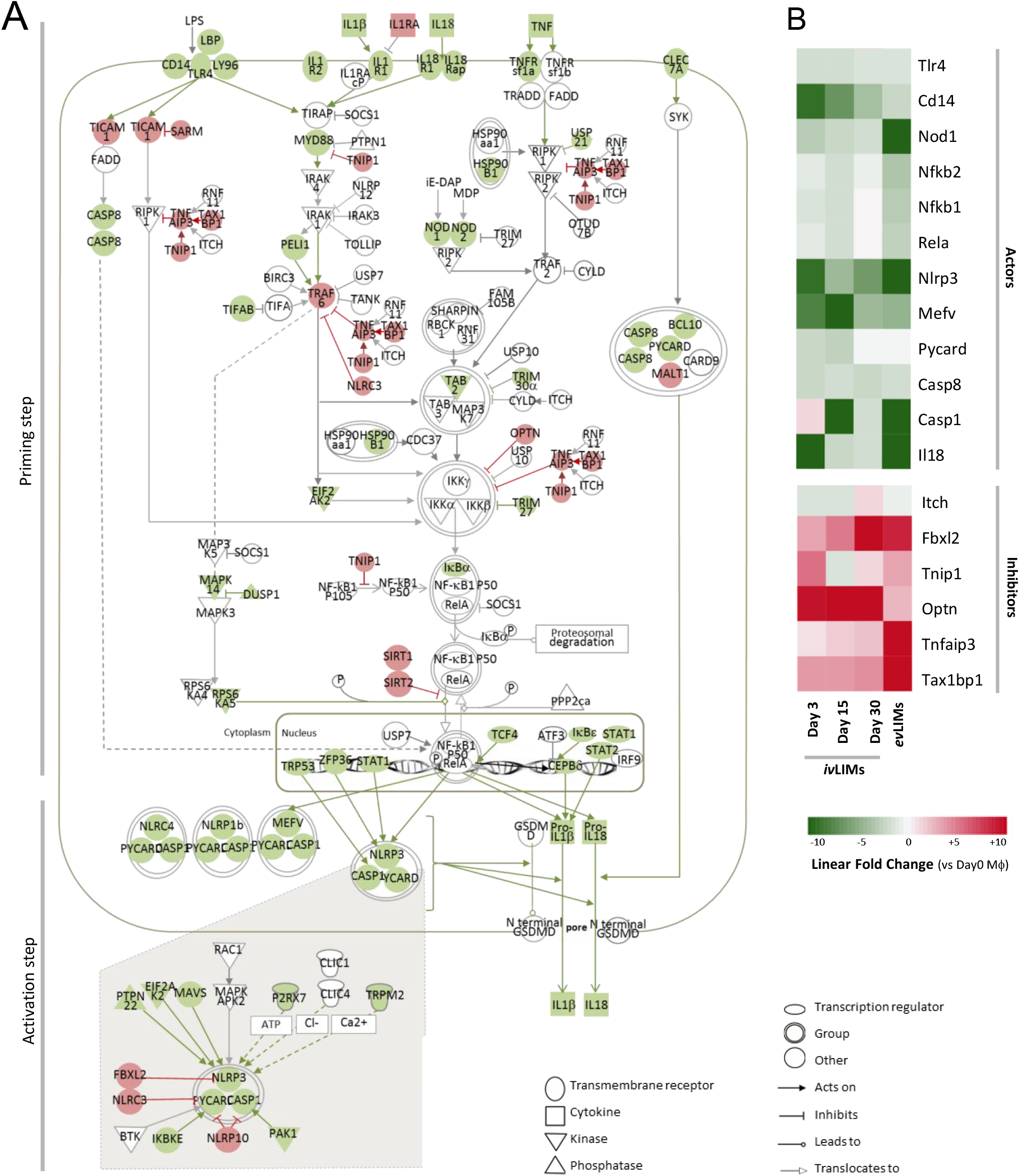
The anti-pyroptotic phenotype of *iv*LIMs and *ev*LIMs correlates with dual transcriptomic repression of the corresponding pathway. **(A)** Subversion of inflammasome priming and activation steps, and pyroptosis in *iv*LIMs at day 3 PI as assessed by RNAseq analysis. Significantly increased (linear FC > +1.25) and decreased (linear FC < −1.25) mRNA abundances are depicted in red and green color, respectively. **(B)** Heatmap of transcriptomic changes observed by RT-qPCR analysis in *iv*LIMs at days 3, 15 and 30 PI (n = 3 independent experiments) and in lesion-derived *ev*LIMs (n = 4 RAG2-KO mice). The analysed transcripts were selected based on our RNAseq results obtained with *iv*LIMs at day 3 PI.

In conclusion, similar to apoptosis and even necroptosis (see Supplementary Figure 4), *Leishmania* dampen the pyroptosis pathway by a unique transcriptomic subversion strategy distinct from other pathogens which interfere with the pyroptotic response by (i) blocking NLRP1, NLRP3 or caspase-1 activation (e.g. Kaposi’s sarcoma-associated herpesvirus, *Mycobacterium tuberculosis;* cowpox virus) (Gregory et al., 2011; Rastogi et al., 2021; Ray et al., 1992), (ii) preventing AIM2 activation by avoiding bacterial DNA release into the macrophage cytosol (*Legionella pneumophila)* (Ge et al., 2012), (iii) inhibition of gasdermin D cleavage (SARS-CoV2) (Ma et al., 2021), or (iv) inhibition of the NF-κB signaling pathway (Hepatitis B virus) (Yu et al., 2017). In the following, we exploited our RNAseq results to investigate the basis of the observed dichotomic regulation of the LIMs RCD pathways by investigating changes in the transcription factor landscape.

### Pan-RCD inhibition in *iv*LIMs correlates with important changes in the host cell transcription factor landscape

We previously reported that *Leishmania* infection stalls Dendritic Cells (DCs) maturation and inflammasome activation by subverting the Transcription Factor (TF) landscape of these cells (Lecoeur et al., 2020b), which we have recently extended to infected BMDMs (Lecoeur et al., 2021). We, therefore, mined our RNAseq for changes in TF expression in LIMs that may explain the anti-RCD phenotype. We revealed significant expression changes in *iv*LIMs at day 3 PI compared to non-infected controls in 88 transcripts for TFs or TF components, most of which were reduced in abundance following infection (Figure 6), including members of the AP1 complex and the NF-kB family previously linked to cell death gene expression (Ameyar et al.; Barkett and Gilmore, 1999). A direct link between the reduced TF abundance and the dichotomic expression changes in LIMs could be established for (i) *stat1* and *trp53* regulating caspase gene expression (Chin et al., 1997; Gupta et al., 2001; Xu et al., 2014), (ii) *trp53* controlling expression of *fa*s, *bbc3, ppmaip1, bcl2l11* and *bax* ((Aubrey et al., 2018), (iii) *ppar* γ, which is increased during infection and inhibits the expression of pro-apoptotic *bax* (Arnett et al., 2018), and iv) *NF-κB* transcription factor family that regulates *birc2, nlrp3, pycard* and *il1β* and is itself inhibited at various levels during *Leishmania* infection (Cornut et al., 2020; Lecoeur et al., 2021; Lecoeur et al., 2020a). Reduced expression of NF-κB family members is likely regulated at epigenetic levels as we have previously shown for *L*.*am*-infected BALB/c macrophages (Lecoeur et al., 2020a). Similarly, links between increased TF abundance and dichotomic expression changes for *pparγ* and its target genes *mcl1* and *bid* (Arnett et al., 2018; Bonofiglio et al., 2011) or *cebpβ* that regulates *tnfaip3* expression (Lai et al., 2013) were also established.

**Figure 6:**
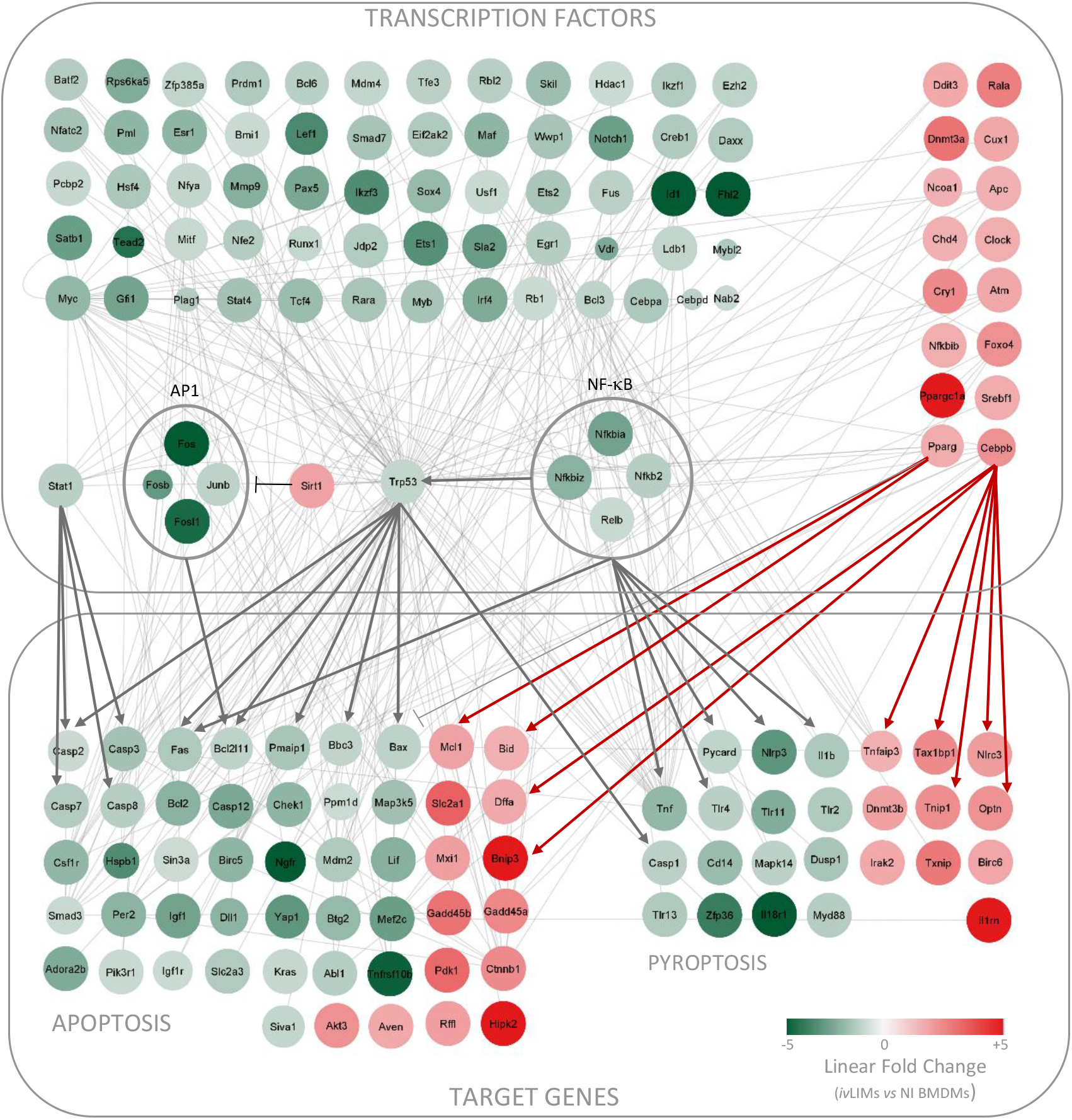
STRING network analysis. Expression changes observed by RNAseq analysis for transcription factors and apoptosis-/pyroptosis-related target genes *iv*LIMs compared to uninfected control at day 3 PI. Significantly up-regulated (linear FC > +1.25) and down-regulated (linear FC < −1.25) genes are depicted in red and green color respectively. *Sta1, nf-κb, trp53, pparγ* and *cebpβ* represent TFs that are known to regulate target genes implicated in the apoptosis and pyroptosis pathways.

In conclusion, *Leishmania* has evolved nuclear subversion strategies that change gene expression control by manipulating the host cell TF landscape, which itself may be regulated by infection-induced changes in epigenetic regulation (Lecoeur et al., 2020a). Transcriptional subversion of RCD pathways seems a common theme during intracellular infection, with various viruses (kaposi virus, Human T Lymphotropic virus-1, Hepatitis viruses), intracellular bacteria (*Rickettsia rickettsii, Legionella pneumophila*) and other parasites (*Toxoplasma gondii*) promoting NF-κB activation to generate an anti-apoptotic profile of their host cell (Clifton et al., 1998; Heussler et al., 2001; Losick et al., 2010; Zamaraev et al., 2020). In contrast, we and others have clearly established that *Leishmania* has evolved multiple strategies to inhibit the TLR/NF-κB and TNFR/NF-κB pathways to avoid a leishmanicidal, pro-inflammatory response (Figure 5) (Lecoeur et al., 2020a). As a result, these parasites were constrained to adopt alternative strategies to interfere with macrophage RCD processes to keep their host cell alive, which proposes *Leishmania* as a powerful new biological tool to probe mechanisms underlying macrophage cell death and longevity.

## Conclusion

Our results uncover a potent and persistent pan-inhibitory effect of intercellular *L. amazonensis* on macrophage RCD *in vitro* and *in vivo* that sustains parasite long-term survival and enables chronic infection. Pleotropic inhibition is regulated at transcriptional levels and consequence of changes in the TF landscape, which cause important expression changes that dampen the pro-RCD and enhance the anti-RCD components specific to apoptosis, pyroptosis and necroptosis, as well as components that are shared between these different forms of cell death. The mechanisms underlying *Leishmania*-mediated subversion of TF-dependent expression control are different from the strategies developed by other pathogens, which include NF-κB-mediated induction of anti-RCD genes or degradation of pro-RCD mediators (Abu-Zant et al., 2007; Robinson and Aw, 2016). Our recent reports on epigenetic and metabolic reprogramming of LIMs (Lecoeur et al., 2020a; Zhang et al., 2022) suggest that changes in TF expression may be regulated by DNA and histone modifications, non-coding RNAs, or changes in host cell glycolytic energy-production and expression of intermediate metabolites that have been linked to RCD resistance (Huang et al., 2015; Huang et al., 2013; Li et al., 2021; Pradelli et al., 2010). Our study opens important new questions on (i) the parasite-released factors that modulate epigenetic, transcriptional and metabolic control in infected LIMs, (ii) the complex interplay between these levels of control in establishing the anti-inflammatory and anti-RCD LIMs phenotype, and (iii) the possibility to design novel host-directed strategy that can rescue the pro-inflammatory and pro-RCD pathways to eliminate intracellular parasites and clear chronic infection.

## ACKNOWLEDGEMENTS

We acknowledge the financial support of the Institut Pasteur (Paris), the Région Ile-de-France (program DIM1Health), the Institut Pasteur International Direction to the International Mixed Unit “Inflammation and *Leishmania* Infection”, the ANR 21CE15004601 ELATION, and the Institut national de la santé et de la recherche médicale (INSERM). Sheng Zhang is part of the Pasteur - Paris University (PPU) International PhD program. This project has received funding from the CNBG company, China.

## SUPPLEMENTARY FIGURE LEGENDS

**Supplementary figure 1: Monitoring of the *L. amazonensis* amastigote load in *iv*LIMs**. C57BL/6 BMDMs were seeded on glass coverslips, infected with mCherry *L. amazonensis* amastigotes. Parasite load was monitored for up to 15 days PI. Parasite nuclei were revealed by Hoechst 33342 staining **(A)**, counted by fluorescence microscopy and the number of parasites per infected macrophage was plotted against time until day 15 PI **(B)**.

**Supplementary figure 2: Assessment of YO-PRO-1 staining in *iv*LIMs and *ev*LIMs. (A)** BMDMs infected with mCherry-transgenic *L. amazonensis* amastigotes (*iv*LIMs) were stained with the nuclear marker Hoechst 33342 (blue) and the cell death marker YO-PRO-1 (green) and monitored microscopically using an EVOS M5000 microscope (20x objective). Note that the single apoptotic cell enlarged in the insert is non-infected (mCherry negative). **(B)** Time course of BMDM (dotted line) and LIMs (continuous line) cell density in culture (one representative experiment is shown out of two). The number of macrophages was determined by counting 5 fields. Projections of the time corresponding to a 50% reduction in macrophage number are indicated for both non-infected (a) and infected (b) cultures. **(C)** Representative picture of *ev*LIMs maintained in complete medium for 15 days post-isolation. Superposition of images of macrophage nuclei (Hoechst 33342 staining), dead macrophages (YO-PRO-1 staining) and parasites (mCherry fluorescence).

**Supplementary figure 3: RNAseq analysis**.

PCA analysis (A) and Volcano plots (B) of expression changes observed non-infected BMDMs (NI) or *L. amazonensis-*infected BMDMs (I) at 3-day PI that were untreated (Ctrl) or treated by LPS (LPS) or LPS and ATP (LPS_ATP) (n = 3 independent biological replicates).

**Supplementary figure 4: Transcriptomic analysis of the anti-necroptosis phenotype in *iv*LIMs and *ev*LIMs**.

**(A)** Subversion of the necroptosis pathway in *iv*LIMs at day 3 PI as assessed by RNAseq analysis. Significantly up-regulated (linear FC > +1.25) and down-regulated (linear FC < −1.25) genes are depicted in red and green color respectively. **(B)** Heatmap of transcriptomic changes observed by RT-qPCR analysis in *iv*LIMs at days 3, 15 and 30 PI (n = 3 independent experiments) and in lesion-derived *ev*LIMs (n = 4 RAG2-KO mice). The analysed transcripts were selected based on our RNAseq results obtained with *iv*LIMs on day 3 PI.

## REFERENCES

Abu-Zant, A., Jones, S., Asare, R., Suttles, J., Price, C., Graham, J., and Kwaik, Y.A. (2007). Anti-apoptotic signalling by the Dot/Icm secretion system of L. pneumophila. Cellular microbiology 9, 246–264.

Akarid, K., Arnoult, D., Micic-Polianski, J., Sif, J., Estaquier, J., and Ameisen, J.C. (2004). Leishmania major-mediated prevention of programmed cell death induction in infected macrophages is associated with the repression of mitochondrial release of cytochrome c. Journal of leukocyte biology 76, 95–103.

Ameyar, M., Wisniewska, M., and Weitzman, J.B. (2003). A role for AP-1 in apoptosis: the case for and against. Biochimie 85, 747–752.

Arango Duque, G., and Descoteaux, A. (2015). Leishmania survival in the macrophage: where the ends justify the means. Current opinion in microbiology 26, 32–40.

Arnett, E., Weaver, A.M., Woodyard, K.C., Montoya, M.J., Li, M., Hoang, K.V., Hayhurst, A., Azad, A.K., and Schlesinger, L.S. (2018). PPARgamma is critical for Mycobacterium tuberculosis induction of Mcl-1 and limitation of human macrophage apoptosis. PLoS pathogens 14, e1007100.

Aubrey, B.J., Kelly, G.L., Janic, A., Herold, M.J., and Strasser, A. (2018). How does p53 induce apoptosis and how does this relate to p53-mediated tumour suppression? Cell death and differentiation 25, 104–113.

Barkett, M., and Gilmore, T.D. (1999). Control of apoptosis by Rel/NF-kappaB transcription factors. Oncogene 18, 6910–6924.

Bedoui, S., Herold, M.J., and Strasser, A. (2020). Emerging connectivity of programmed cell death pathways and its physiological implications. Nature reviews Molecular cell biology 21, 678–695.

Behar, S.M., Martin, C.J., Booty, M.G., Nishimura, T., Zhao, X., Gan, H.X., Divangahi, M., and Remold, H.G. (2011). Apoptosis is an innate defense function of macrophages against Mycobacterium tuberculosis. Mucosal immunology 4, 279–287.

Bergsbaken, T., Fink, S.L., and Cookson, B.T. (2009). Pyroptosis: host cell death and inflammation. Nature reviews Microbiology 7, 99–109.

Birge, R.B., and Ucker, D.S. (2008). Innate apoptotic immunity: the calming touch of death. Cell death and differentiation 15, 1096–1102.

Bonofiglio, D., Cione, E., Vizza, D., Perri, M., Pingitore, A., Qi, H., Catalano, S., Rovito, D., Genchi, G., and Ando, S. (2011). Bid as a potential target of apoptotic effects exerted by low doses of PPARgamma and RXR ligands in breast cancer cells. Cell cycle 10, 2344–2354.

Brune, W., and Andoniou, C.E. (2017). Die Another Day: Inhibition of Cell Death Pathways by Cytomegalovirus. Viruses 9.

Chau, B.N., Cheng, E.H., Kerr, D.A., and Hardwick, J.M. (2000). Aven, a novel inhibitor of caspase activation, binds Bcl-xL and Apaf-1. Molecular cell 6, 31–40.

Chin, Y.E., Kitagawa, M., Kuida, K., Flavell, R.A., and Fu, X.Y. (1997). Activation of the STAT signaling pathway can cause expression of caspase 1 and apoptosis. Molecular and cellular biology 17, 5328–5337.

Chow, S.H., Deo, P., and Naderer, T. (2016). Macrophage cell death in microbial infections. Cellular microbiology 18, 466–474.

Christgen, S., Place, D.E., and Kanneganti, T.D. (2020). Toward targeting inflammasomes: insights into their regulation and activation. Cell research 30, 315–327.

Cianciulli, A., Porro, C., Calvello, R., Trotta, T., and Panaro, M.A. (2018). Resistance to apoptosis in Leishmania infantum-infected human macrophages: a critical role for anti-apoptotic Bcl-2 protein and cellular IAP1/2. Clinical and experimental medicine 18, 251–261.

Clifton, D.R., Goss, R.A., Sahni, S.K., van Antwerp, D., Baggs, R.B., Marder, V.J., Silverman, D.J., and Sporn, L.A. (1998). NF-kappa B-dependent inhibition of apoptosis is essential for host cellsurvival during Rickettsia rickettsii infection. Proceedings of the National Academy of Sciences of the United States of America 95, 4646–4651.

Cordsmeier, A., Wagner, N., Luhrmann, A., and Berens, C. (2019). Defying Death - How Coxiella burnetii Copes with Intentional Host Cell Suicide. The Yale journal of biology and medicine 92, 619–628.

Cornut, M., Bourdonnay, E., and Henry, T. (2020). Transcriptional Regulation of Inflammasomes. International journal of molecular sciences 21.

Dadsena, S., Jenner, A., and Garcia-Saez, A.J. (2021). Mitochondrial outer membrane permeabilization at the single molecule level. Cellular and molecular life sciences : CMLS 78, 3777–3790.

Darzynkiewicz, Z., Holden, E., Orfao, A., Telford, W.G., and Wlodkowic, D. (2011). Recent advances in cytometry, Part B, Vol 103, Fifth edition edn (Elsevier).

Dashzeveg, N., and Yoshida, K. (2015). Cell death decision by p53 via control of the mitochondrial membrane. Cancer letters 367, 108–112.

Dobin, A., Davis, C.A., Schlesinger, F., Drenkow, J., Zaleski, C., Jha, S., Batut, P., Chaisson, M., and Gingeras, T.R. (2013). STAR: ultrafast universal RNA-seq aligner. Bioinformatics 29, 15–21.

Donovan, M.J., Maciuba, B.Z., Mahan, C.E., and McDowell, M.A. (2009). Leishmania infection inhibits cycloheximide-induced macrophage apoptosis in a strain-dependent manner. Experimental parasitology 123, 58–64.

Ferreira, C., Estaquier, J., and Silvestre, R. (2021). Immune-metabolic interactions between Leishmania and macrophage host. Current opinion in microbiology 63, 231–237.

Frank, D., and Vince, J.E. (2019). Pyroptosis versus necroptosis: similarities, differences, and crosstalk. Cell death and differentiation 26, 99–114.

Galluzzi, L., Vitale, I., Aaronson, S.A., Abrams, J.M., Adam, D., Agostinis, P., Alnemri, E.S., Altucci, L., Amelio, I., Andrews, D.W., et al. (2018). Molecular mechanisms of cell death: recommendations of the Nomenclature Committee on Cell Death 2018. Cell death and differentiation 25, 486–541.

Gatto, M., Borim, P.A., Wolf, I.R., Fukuta da Cruz, T., Ferreira Mota, G.A., Marques Braz, A.M., Casella Amorim, B., Targino Valente, G., de Assis Golim, M., Venturini, J., et al. (2020). Transcriptional analysis of THP-1 cells infected with Leishmania infantum indicates no activation of the inflammasome platform. PLoS neglected tropical diseases 14, e0007949.

Ge, J., Gong, Y.N., Xu, Y., and Shao, F. (2012). Preventing bacterial DNA release and absent in melanoma 2 inflammasome activation by a Legionella effector functioning in membrane trafficking. Proceedings of the National Academy of Sciences of the United States of America 109, 6193–6198.

Giri, J., Srivastav, S., Basu, M., Palit, S., Gupta, P., and Ukil, A. (2016). Leishmania donovani Exploits Myeloid Cell Leukemia 1 (MCL-1) Protein to Prevent Mitochondria-dependent Host Cell Apoptosis. The Journal of biological chemistry 291, 3496–3507.

Gregory, S.M., Davis, B.K., West, J.A., Taxman, D.J., Matsuzawa, S., Reed, J.C., Ting, J.P., and Damania, B. (2011). Discovery of a viral NLR homolog that inhibits the inflammasome. Science 331, 330–334.

Grimsley, C., and Ravichandran, K.S. (2003). Cues for apoptotic cell engulfment: eat-me, don’t eat-me and come-get-me signals. Trends in cell biology 13, 648–656.

Grootjans, S., Vanden Berghe, T., and Vandenabeele, P. (2017). Initiation and execution mechanisms of necroptosis: an overview. Cell death and differentiation 24, 1184–1195.

Gupta, S., Radha, V., Furukawa, Y., and Swarup, G. (2001). Direct transcriptional activation of human caspase-1 by tumor suppressor p53. The Journal of biological chemistry 276, 10585–10588.

Hacker, G. (2018). Apoptosis in infection. Microbes and infection 20, 552–559.

Han, H., Cho, J.W., Lee, S., Yun, A., Kim, H., Bae, D., Yang, S., Kim, C.Y., Lee, M., Kim, E., et al. (2018). TRRUST v2: an expanded reference database of human and mouse transcriptional regulatory interactions. Nucleic acids research 46, D380–D386.

Haneklaus, M., O’Neil, J.D., Clark, A.R., Masters, S.L., and O’Neill, L.A.J. (2017). The RNA-binding protein Tristetraprolin (TTP) is a critical negative regulator of the NLRP3 inflammasome. The Journal of biological chemistry 292, 6869–6881.

Heilig, R., Dick, M.S., Sborgi, L., Meunier, E., Hiller, S., and Broz, P. (2018). The Gasdermin-D pore acts as a conduit for IL-1beta secretion in mice. European journal of immunology 48, 584–592.

Heussler, V.T., Kuenzi, P., and Rottenberg, S. (2001). Inhibition of apoptosis by intracellular protozoan parasites. International journal for parasitology 31, 1166–1176.

Holler, N., Zaru, R., Micheau, O., Thome, M., Attinger, A., Valitutti, S., Bodmer, J.L., Schneider, P., Seed, B., and Tschopp, J. (2000). Fas triggers an alternative, caspase-8-independent cell death pathway using the kinase RIP as effector molecule. Nature immunology 1, 489–495.

Hu, X.M., Li, Z.X., Lin, R.H., Shan, J.Q., Yu, Q.W., Wang, R.X., Liao, L.S., Yan, W.T., Wang, Z., Shang, L., et al. (2021). Guidelines for Regulated Cell Death Assays: A Systematic Summary, A Categorical Comparison, A Prospective. Frontiers in cell and developmental biology 9, 634690.

Huang, C., Sheng, S., Li, R., Sun, X., Liu, J., and Huang, G. (2015). Lactate promotes resistance to glucose starvation via upregulation of Bcl-2 mediated by mTOR activation. Oncology reports 33, 875–884.

Huang, C.Y., Kuo, W.T., Huang, Y.C., Lee, T.C., and Yu, L.C. (2013). Resistance to hypoxia-induced necroptosis is conferred by glycolytic pyruvate scavenging of mitochondrial superoxide in colorectal cancer cells. Cell death & disease 4, e622.

Idziorek, T., Estaquier, J., De Bels, F., and Ameisen, J.C. (1995). YOPRO-1 permits cytofluorometric analysis of programmed cell death (apoptosis) without interfering with cell viability. Journal of immunological methods 185, 249–258.

Jorgensen, I., Rayamajhi, M., and Miao, E.A. (2017). Programmed cell death as a defence against infection. Nature reviews Immunology 17, 151–164.

Kelley, N., Jeltema, D., Duan, Y., and He, Y. (2019). The NLRP3 Inflammasome: An Overview of Mechanisms of Activation and Regulation. International journal of molecular sciences 20.

Kloehn, J., Saunders, E.C., O’Callaghan, S., Dagley, M.J., and McConville, M.J. (2015). Characterization of metabolically quiescent Leishmania parasites in murine lesions using heavy water labeling. PLoS pathogens 11, e1004683.

Kolbrink, B., Riebeling, T., Kunzendorf, U., and Krautwald, S. (2020). Plasma Membrane Pores Drive Inflammatory Cell Death. Frontiers in cell and developmental biology 8, 817.

Kourtzelis, I., Hajishengallis, G., and Chavakis, T. (2020). Phagocytosis of Apoptotic Cells in Resolution of Inflammation. Frontiers in immunology 11, 553.

Kutuk, O., Temel, S.G., Tolunay, S., and Basaga, H. (2010). Aven blocks DNA damage-induced apoptosis by stabilising Bcl-xL. European journal of cancer 46, 2494–2505.

Lai, T.Y., Wu, S.D., Tsai, M.H., Chuang, E.Y., Chuang, L.L., Hsu, L.C., and Lai, L.C. (2013). Transcription of Tnfaip3 is regulated by NF-kappaB and p38 via C/EBPbeta in activated macrophages. PloS one 8, e73153.

Lecoeur, H., de La Llave, E., Osorio, Y.F.J., Goyard, S., Kiefer-Biasizzo, H., Balazuc, A.M., Milon, G., Prina, E., and Lang, T. (2010). Sorting of Leishmania-bearing dendritic cells reveals subtle parasite-induced modulation of host-cell gene expression. Microbes and infection 12, 46–54.

Lecoeur, H., Prina, E., Gutierrez-Sanchez, M., and Spath, G.F. (2021). Going ballistic: Leishmania nuclear subversion of host cell plasticity. Trends in parasitology.

Lecoeur, H., Prina, E., Rosazza, T., Kokou, K., N’Diaye, P., Aulner, N., Varet, H., Bussotti, G., Xing, Y., Milon, G., et al. (2020a). Targeting Macrophage Histone H3 Modification as a Leishmania Strategy to Dampen the NF-kappaB/NLRP3-Mediated Inflammatory Response. Cell Rep 30, 1870–1882 e1874.

Lecoeur, H., Rosazza, T., Kokou, K., Varet, H., Coppee, J.Y., Lari, A., Commere, P.H., Weil, R., Meng, G., Milon, G., et al. (2020b). Leishmania amazonensis Subverts the Transcription Factor Landscape in Dendritic Cells to Avoid Inflammasome Activation and Stall Maturation. Frontiers in immunology 11, 1098.

Li, L.Y., Nie, Z.Y., Zhang, X.Y., Luo, J.M., Yang, L., Wang, Q., and Wang, X.Z. (2021). [The Effect of si-PKM2 on Proliferation and Apoptosis of Acute Leukemic Cells and Its Molecular Mechanism]. Zhongguo shi yan xue ye xue za zhi 29, 1394–1402.

Liao, Y., Smyth, G.K., and Shi, W. (2014). featureCounts: an efficient general purpose program for assigning sequence reads to genomic features. Bioinformatics 30, 923–930.

Lisi, S., Sisto, M., Acquafredda, A., Spinelli, R., Schiavone, M., Mitolo, V., Brandonisio, O., and Panaro, M. (2005). Infection with Leishmania infantum Inhibits actinomycin D-induced apoptosis of human monocytic cell line U-937. The Journal of eukaryotic microbiology 52, 211–217.

Liu, D., and Uzonna, J.E. (2012). The early interaction of Leishmania with macrophages and dendritic cells and its influence on the host immune response. Frontiers in cellular and infection microbiology 2, 83.

Liu, X.F., Xiang, L., Zhou, Q., Carralot, J.P., Prunotto, M., Niederfellner, G., and Pastan, I. (2016). Actinomycin D enhances killing of cancer cells by immunotoxin RG7787 through activation of the extrinsic pathway of apoptosis. Proceedings of the National Academy of Sciences of the United States of America 113, 10666–10671.

Losick, V.P., Haenssler, E., Moy, M.Y., and Isberg, R.R. (2010). LnaB: a Legionella pneumophila activator of NF-kappaB. Cellular microbiology 12, 1083–1097.

Luz, N.F., Khouri, R., Van Weyenbergh, J., Zanette, D.L., Fiuza, P.P., Noronha, A., Barral, A., Boaventura, V.S., Prates, D.B., Chan, F.K., et al. (2018). Leishmania braziliensis Subverts Necroptosis by Modulating RIPK3 Expression. Frontiers in microbiology 9, 2283.

Ma, J., Zhu, F., Zhao, M., Shao, F., Yu, D., Ma, J., Zhang, X., Li, W., Qian, Y., Zhang, Y., et al. (2021). SARS-CoV-2 nucleocapsid suppresses host pyroptosis by blocking Gasdermin D cleavage. The EMBO journal 40, e108249.

Medina, C.B., Mehrotra, P., Arandjelovic, S., Perry, J.S.A., Guo, Y., Morioka, S., Barron, B., Walk, S.F., Ghesquiere, B., Krupnick, A.S., et al. (2020). Metabolites released from apoptotic cells act as tissue messengers. Nature 580, 130–135.

Merkel, O., Wacht, N., Sifft, E., Melchardt, T., Hamacher, F., Kocher, T., Denk, U., Hofbauer, J.P., Egle, A., Scheideler, M., et al. (2012). Actinomycin D induces p53-independent cell death and prolongs survival in high-risk chronic lymphocytic leukemia. Leukemia 26, 2508–2516.

Mocarski, E.S., Upton, J.W., and Kaiser, W.J. (2011). Viral infection and the evolution of caspase 8-regulated apoptotic and necrotic death pathways. Nature reviews Immunology 12, 79–88.

Moore, K.J., Turco, S.J., and Matlashewski, G. (1994). Leishmania donovani infection enhances macrophage viability in the absence of exogenous growth factor. Journal of leukocyte biology 55, 91–98.

Murao, A., Aziz, M., Wang, H., Brenner, M., and Wang, P. (2021). Release mechanisms of major DAMPs. Apoptosis : an international journal on programmed cell death 26, 152–162.

Paik, S., Kim, J.K., Silwal, P., Sasakawa, C., and Jo, E.K. (2021). An update on the regulatory mechanisms of NLRP3 inflammasome activation. Cellular & molecular immunology 18, 1141–1160.

Pandey, R.K., Mehrotra, S., Sharma, S., Gudde, R.S., Sundar, S., and Shaha, C. (2016). Leishmania donovani-Induced Increase in Macrophage Bcl-2 Favors Parasite Survival. Frontiers in immunology 7, 456.

Pasparakis, M., and Vandenabeele, P. (2015). Necroptosis and its role in inflammation. Nature 517, 311–320.

Pfaffl, M.W., Horgan, G.W., and Dempfle, L. (2002). Relative expression software tool (REST) for group-wise comparison and statistical analysis of relative expression results in real-time PCR. Nucleic acids research 30, e36.

Pirbhai, M., Dong, F., Zhong, Y., Pan, K.Z., and Zhong, G. (2006). The secreted protease factor CPAF is responsible for degrading pro-apoptotic BH3-only proteins in Chlamydia trachomatis-infected cells. The Journal of biological chemistry 281, 31495–31501.

Place, D.E., Lee, S., and Kanneganti, T.D. (2021). PANoptosis in microbial infection. Current opinion in microbiology 59, 42–49.

Pradelli, L.A., Beneteau, M., Chauvin, C., Jacquin, M.A., Marchetti, S., Munoz-Pinedo, C., Auberger, P., Pende, M., and Ricci, J.E. (2010). Glycolysis inhibition sensitizes tumor cells to death receptors-induced apoptosis by AMP kinase activation leading to Mcl-1 block in translation. Oncogene 29, 1641–1652.

Priem, D., van Loo, G., and Bertrand, M.J.M. (2020). A20 and Cell Death-driven Inflammation. Trends Immunol 41, 421–435.

Rastogi, S., Ellinwood, S., Augenstreich, J., Mayer-Barber, K.D., and Briken, V. (2021). Mycobacterium tuberculosis inhibits the NLRP3 inflammasome activation via its phosphokinase PknF. PLoS pathogens 17, e1009712.

Ray, C.A., Black, R.A., Kronheim, S.R., Greenstreet, T.A., Sleath, P.R., Salvesen, G.S., and Pickup, D.J. (1992). Viral inhibition of inflammation: cowpox virus encodes an inhibitor of the interleukin-1 beta converting enzyme. Cell 69, 597–604.

Riera Romo, M. (2021). Cell death as part of innate immunity: Cause or consequence? Immunology 163, 399–415.

Robinson, K.S., and Aw, R. (2016). The Commonalities in Bacterial Effector Inhibition of Apoptosis. Trends in microbiology 24, 665–680.

Robinson, N., Ganesan, R., Hegedus, C., Kovacs, K., Kufer, T.A., and Virag, L. (2019). Programmed necrotic cell death of macrophages: Focus on pyroptosis, necroptosis, and parthanatos. Redox biology 26, 101239.

Rodriguez-Gonzalez, J., Wilkins-Rodriguez, A., Argueta-Donohue, J., Aguirre-Garcia, M., and Gutierrez-Kobeh, L. (2016). Leishmania mexicana promastigotes down regulate JNK and p-38 MAPK activation: Role in the inhibition of camptothecin-induced apoptosis of monocyte-derived dendritic cells. Experimental parasitology 163, 57–67.

Rodriguez-Gonzalez, J., Wilkins-Rodriguez, A.A., and Gutierrez-Kobeh, L. (2022). Involvement of Akt and the antiapoptotic protein Bcl-xL in the inhibition of apoptosis of dendritic cells by Leishmania mexicana. Parasite immunology 44, e12917.

Rosazza, T., Lecoeur, H., Blisnick, T., Moya-Nilges, M., Pescher, P., Bastin, P., Prina, E., and Spath, G.F. (2020). Dynamic imaging reveals surface exposure of virulent Leishmania amastigotes during pyroptosis of infected macrophages. Journal of cell science 134.

Roy, S., Gupta, P., Palit, S., Basu, M., Ukil, A., and Das, P.K. (2017). The role of PD-1 in regulation of macrophage apoptosis and its subversion by Leishmania donovani. Clinical & translational immunology 6, e137.

Ruhland, A., Leal, N., and Kima, P.E. (2007). Leishmania promastigotes activate PI3K/Akt signalling to confer host cell resistance to apoptosis. Cellular microbiology 9, 84–96.

Saha, G., Khamar, B.M., Prerna, K., Kumar, M., and Dubey, V.K. (2019a). BLIMP-1 Plays Important Role in the Regulation of Macrophage Pyroptosis for the Growth and Multiplication of Leishmania donovani. ACS infectious diseases 5, 2087–2095.

Saha, G., Khamar, B.M., Singh, O.P., Sundar, S., and Dubey, V.K. (2019b). Leishmania donovani evades Caspase 1 dependent host defense mechanism during infection. International journal of biological macromolecules 126, 392–401.

Saunders, E.C., and McConville, M.J. (2020). Immunometabolism of Leishmania granulomas. Immunology and cell biology 98, 832–844.

Shinkai, Y., Rathbun, G., Lam, K.P., Oltz, E.M., Stewart, V., Mendelsohn, M., Charron, J., Datta, M., Young, F., Stall, A.M., et al. (1992). RAG-2-deficient mice lack mature lymphocytes owing to inability to initiate V(D)J rearrangement. Cell 68, 855–867.

Swanson, K.V., Deng, M., and Ting, J.P. (2019). The NLRP3 inflammasome: molecular activation and regulation to therapeutics. Nature reviews Immunology 19, 477–489.

Tang, D., Kang, R., Berghe, T.V., Vandenabeele, P., and Kroemer, G. (2019). The molecular machinery of regulated cell death. Cell research 29, 347–364.

Tummers, B., and Green, D.R. (2022). The Evolution of Regulated Cell Death Pathways in Animals and Their Evasion by Pathogens. Physiol Rev 102.

Van Assche, T., Deschacht, M., da Luz, R.A., Maes, L., and Cos, P. (2011). Leishmania-macrophage interactions: insights into the redox biology. Free radical biology & medicine 51, 337–351.

Voth, D.E., and Heinzen, R.A. (2009). Sustained activation of Akt and Erk1/2 is required for Coxiella burnetii antiapoptotic activity. Infection and immunity 77, 205–213.

Weiss, G., and Schaible, U.E. (2015). Macrophage defense mechanisms against intracellular bacteria. Immunological reviews 264, 182–203.

Whitfield, J., Neame, S.J., Paquet, L., Bernard, O., and Ham, J. (2001). Dominant-negative c-Jun promotes neuronal survival by reducing BIM expression and inhibiting mitochondrial cytochrome c release. Neuron 29, 629–643.

Xia, X., Wang, X., Zheng, Y., Jiang, J., and Hu, J. (2019). What role does pyroptosis play in microbial infection? Journal of cellular physiology 234, 7885–7892.

Xia, X.J., Lei, L.C., Wang, S., Hu, J.H., and Zhang, G.P. (2020). Necroptosis and its role in infectious diseases. Apoptosis : an international journal on programmed cell death 25, 169–178.

Xu, X., Wen, H., Hu, Y., Yu, H., Zhang, Y., Chen, C., and Pan, X. (2014). STAT1-caspase 3 pathway in the apoptotic process associated with steroid-induced necrosis of the femoral head. Journal of molecular histology 45, 473–485.

Yu, P., Zhang, X., Liu, N., Tang, L., Peng, C., and Chen, X. (2021). Pyroptosis: mechanisms and diseases. Signal transduction and targeted therapy 6, 128.

Yu, X., Lan, P., Hou, X., Han, Q., Lu, N., Li, T., Jiao, C., Zhang, J., Zhang, C., and Tian, Z. (2017). HBV inhibits LPS-induced NLRP3 inflammasome activation and IL-1beta production via suppressing the NF-kappaB pathway and ROS production. Journal of hepatology 66, 693–702.

Zamaraev, A.V., Zhivotovsky, B., and Kopeina, G.S. (2020). Viral Infections: Negative Regulators of Apoptosis and Oncogenic Factors. Biochemistry Biokhimiia 85, 1191–1201.

Zhang, S., Lecoeur, H., Varet, H., Legendre, R., Mahtal, N., Proux, C., Aulner, N., Shorte, S., Granjean, C., Bousso, P., et al. (2022). Immunometabolic profiling of in vitro and ex vivo Leishmania-infected macrophages (LIMs) reveals unique polarization and bioenergetic signatures. BioRxiv, DOI: 101101/20220908507100.

