## Supplemental figures for "*Leishmania amazonensis* controls macrophage-regulated cell death to establish chronic infection *in vitro* and *in vivo*"

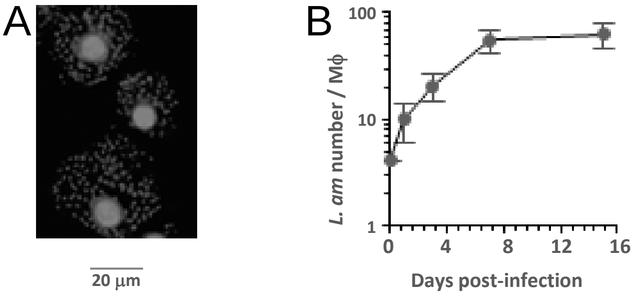


Supplementary Figure1 , Lecoeur H. et al.

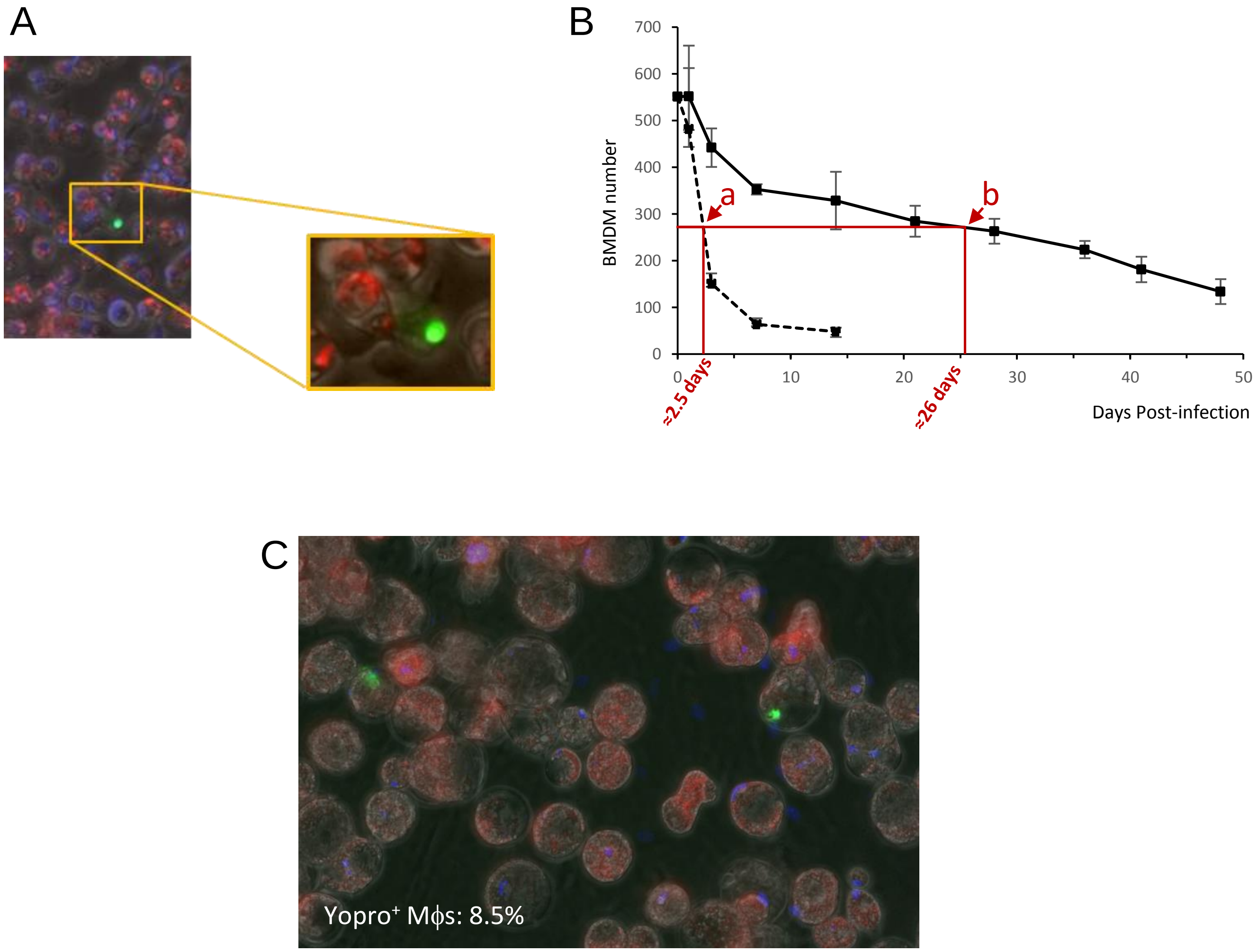


Supplementary Figure 2 , Lecoeur H. et al.

A

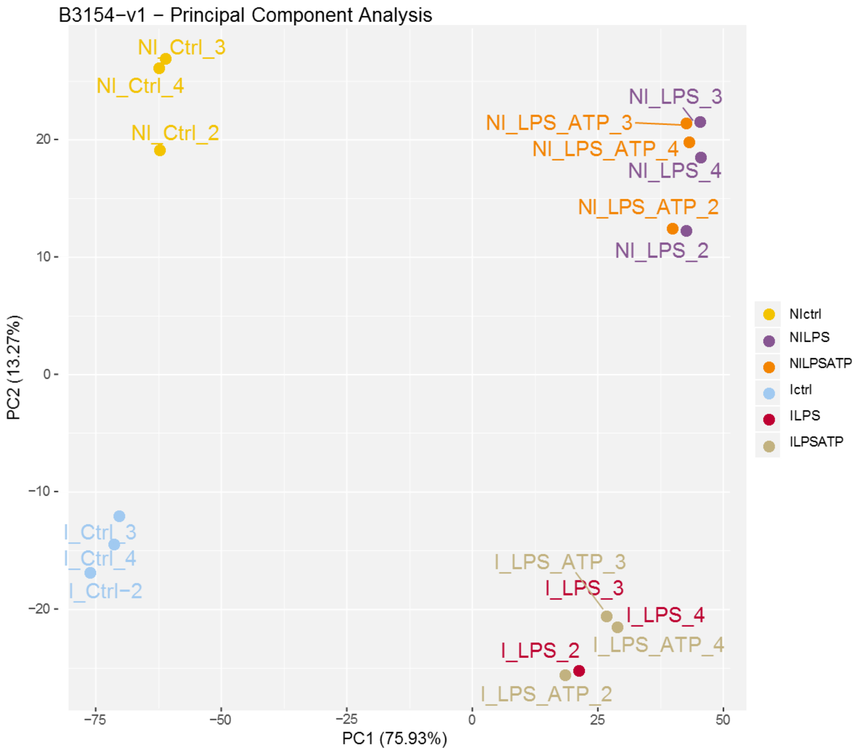


B

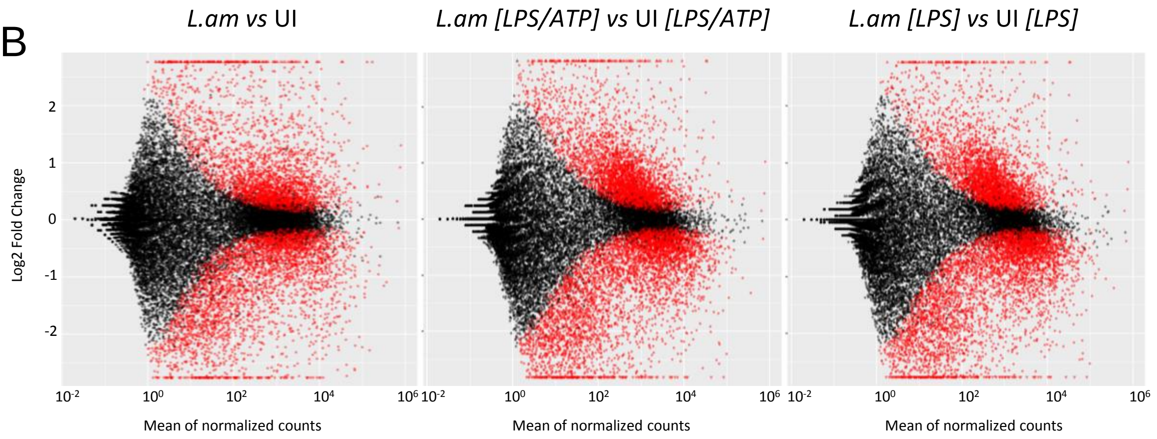


Supplementary Figure 3 , Lecoeur H. et al.

A 1

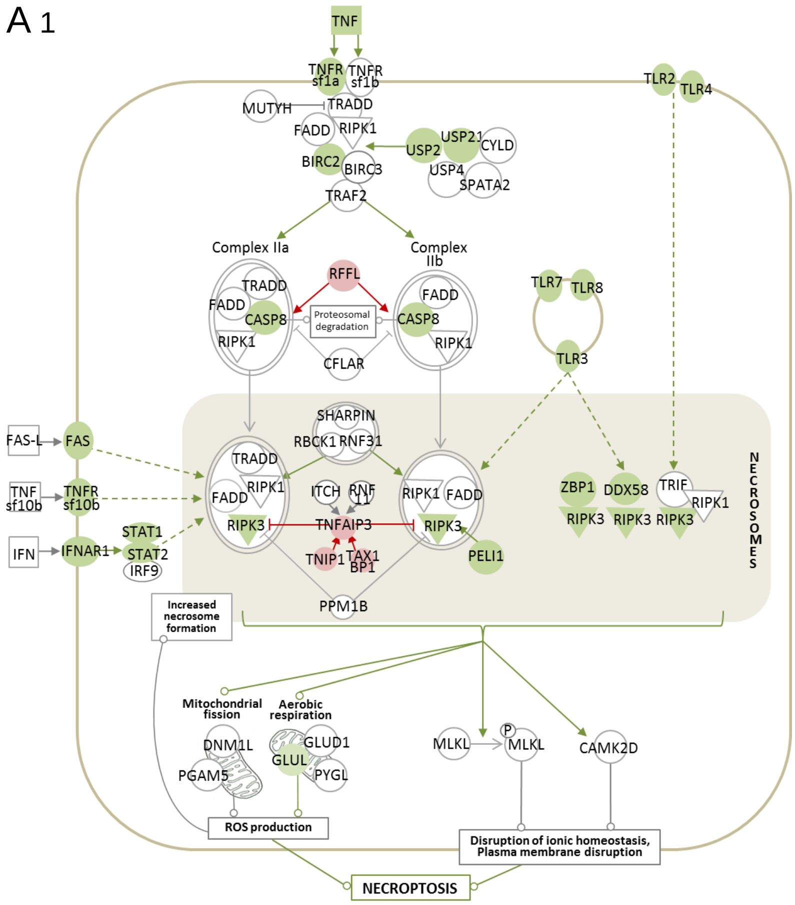


2

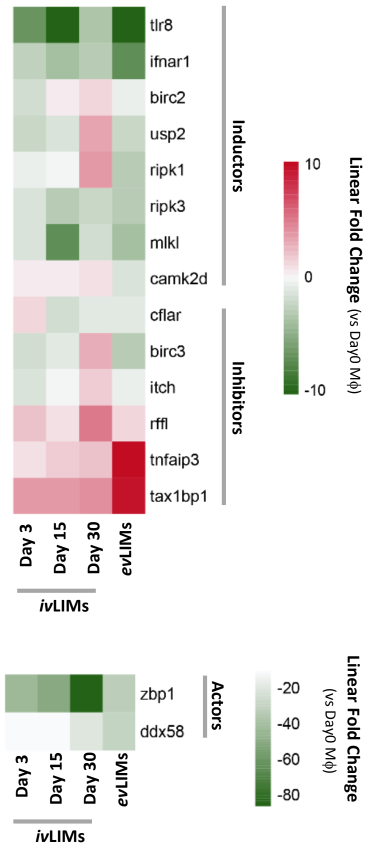


Supplementary Figure 4, Lecoeur H. et al.
