## Supplementary Table 1 for "*Leishmania amazonensis* controls macrophage-regulated cell death to establish chronic infection *in vitro* and *in vivo*"

| Id | gene_name | Necroptosis activator 0/ inhibitor 1 |
|---|---|---|
| ENSMUSG00000057367 | **Birc2** | 1 |
| ENSMUSG00000032000 | **birc3** | 1 |
| ENSMUSG00000026278 | **bok** | 1 |
| ENSMUSG00000026029 | **Casp8** | 1 |
| ENSMUSG00000026031 | **Cflar** | 1 |
| ENSMUSG00000040314 | **ctsg** | 0 |
| ENSMUSG00000036712 | **Cyld** | 0 |
| ENSMUSG00000040296 | **Ddx58** | 0 |
| ENSMUSG00000031077 | **fadd** | 0 |
| ENSMUSG00000024778 | **Fas** | 0 |
| ENSMUSG00000000817 | **fasl** | 0 |
| ENSMUSG00000026473 | **Glul** | 0 |
| ENSMUSG00000022967 | **Ifnar1** | 0 |
| ENSMUSG00000022971 | **Ifnar2** | 0 |
| ENSMUSG00000060733 | **ipmk** | 0 |
| ENSMUSG00000002325 | **Irf9** | 0 |
| ENSMUSG00000027598 | **itch** | 1 |
| ENSMUSG00000028284 | **map3k7** | 1 |
| ENSMUSG00000037523 | **Mavs** | 0 |
| ENSMUSG00000012519 | **Mlkl** | 0 |
| ENSMUSG00000028687 | **mutyh** | 1 |
| ENSMUSG00000020134 | **Peli1** | 0 |
| ENSMUSG00000029500 | **pgam5** | 0 |
| ENSMUSG00000021868 | **ppif** | 0 |
| ENSMUSG00000061130 | **ppm1b** | 1 |
| ENSMUSG00000021069 | **pygl** | 0 |
| ENSMUSG00000027466 | **rbck1** | 1 |
| ENSMUSG00000021408 | **Ripk1** | 0 |
| ENSMUSG00000022221 | **Ripk3** | 0 |
| ENSMUSG00000028557 | **rnf11** | 1 |
| ENSMUSG00000047030 | **spata2** | 0 |
| ENSMUSG00000026104 | **Stat1** | 0 |
| ENSMUSG00000040033 | **Stat2** | 0 |
| ENSMUSG00000004535 | **Tax1bp1** | 1 |
| ENSMUSG00000047123 | **Ticam1** | 0 |
| ENSMUSG00000027995 | **Tlr2** | 0 |
| ENSMUSG00000031639 | **tlr3** | 0 |
| ENSMUSG00000039005 | **Tlr4** | 0 |
| ENSMUSG00000044583 | **Tlr7** | 0 |
| ENSMUSG00000040522 | **Tlr8** | 0 |
| ENSMUSG00000019850 | **Tnfaip3** | 1 |
| ENSMUSG00000022074 | **Tnfrsf10b** | 0 |
| ENSMUSG00000037613 | **tnfrsf23** | 0 |
| ENSMUSG00000039304 | **Tnfsf10** | 0 |
| ENSMUSG00000020400 | **Tnip1** | 1 |
| ENSMUSG00000031887 | **Tradd** | 0 |
| ENSMUSG00000032010 | **Usp2** | 0 |
| ENSMUSG00000053483 | **Usp21** | 0 |
| ENSMUSG00000032612 | **Usp4** | 0 |

| Id | gene_name | Necroptosis activator 0/ inhibitor 1 |
|---|---|---|
| ENSMUSG00000030189 | **ybx3** | 1 |
| ENSMUSG00000027514 | **Zbp1** | 0 |
| ENSMUSG00000022552 | **Sharpin** | 0 |
| ENSMUSG00000027466 | **rbck1** | 0 |
| ENSMUSG00000047098 | **rnf31** | 0 |
| ENSMUSG00000024401 | **Tnf** | 0 |
| ENSMUSG00000030341 | **Tnfrsf1a** | 0 |
| ENSMUSG00000028599 | **Tnfrsf1b** | 0 |
| ENSMUSG00000020696 | **Rffl** | 0 |
| ENSMUSG00000027514 | **Zbp1** | 0 |
| ENSMUSG00000053819 | **Camk2d** | 0 |
| ENSMUSG00000022789 | **Dnm1l** | 0 |
| ENSMUSG00000021794 | **glud1** | 0 |

| Id | gene_name | Pyroptosis activator 0/ inhibitor 1 |
|---|---|---|
| ENSMUSG00000037860 | Aim2 | 0 |
| ENSMUSG00000010911 | apip | 1 |
| ENSMUSG00000026628 | Atf3 | 0 |
| ENSMUSG00000028191 | Bcl10 | 0 |
| ENSMUSG00000032000 | Birc3 | 0 |
| ENSMUSG00000031201 | brcc3 | 0 |
| ENSMUSG00000031264 | Btk | 0 |
| ENSMUSG00000026928 | card9 | 0 |
| ENSMUSG00000025888 | Casp1 | 0 |
| ENSMUSG00000026029 | Casp8 | 0 |
| ENSMUSG00000051439 | Cd14 | 0 |
| ENSMUSG00000019471 | cdc37 | 0 |
| ENSMUSG00000071637 | Cebpd | 0 |
| ENSMUSG00000079293 | Clec7a | 0 |
| ENSMUSG00000007041 | Clic1 | 0 |
| ENSMUSG00000037242 | Clic4 | 0 |
| ENSMUSG00000020638 | Cmpk2 | 0 |
| ENSMUSG00000021939 | Ctsb | 0 |
| ENSMUSG00000036712 | Cyld | 1 |
| ENSMUSG00000004099 | Dnmt1 | ND |
| ENSMUSG00000027478 | Dnmt3b | 1 |
| ENSMUSG00000024190 | Dusp1 | 1 |
| ENSMUSG00000024079 | Eif2ak2 | 0 |
| ENSMUSG00000031077 | fadd | 0 |
| ENSMUSG00000046034 | otulin | 1 |
| ENSMUSG00000032507 | Fbxl2 | 1 |
| ENSMUSG00000022575 | gsdmd | 0 |
| ENSMUSG00000021270 | Hsp90aa1 | 0 |
| ENSMUSG00000020048 | Hsp90b1 | 0 |
| ENSMUSG00000031537 | ikbkb | 0 |
| ENSMUSG00000042349 | Ikbke | 0 |
| ENSMUSG00000004221 | Ikbkg | 1 |
| ENSMUSG00000039217 | Il18 | 0 |
| ENSMUSG00000070427 | Il18bp | 0 |
| ENSMUSG00000026070 | Il18r1 | 0 |
| ENSMUSG00000026068 | Il18rap | 0 |
| ENSMUSG00000027398 | Il1b | 0 |
| ENSMUSG00000026072 | Il1r1 | 0 |
| ENSMUSG00000026073 | Il1r2 | 1 |
| ENSMUSG00000022514 | Il1rap | 0 |
| ENSMUSG00000026981 | Il1rn | 1 |
| ENSMUSG00000031392 | Irak1 | 0 |
| ENSMUSG00000060477 | Irak2 | 0 |
| ENSMUSG00000020227 | irak3 | 1 |
| ENSMUSG00000059883 | Irak4 | 0 |
| ENSMUSG00000002325 | Irf9 | 0 |
| ENSMUSG00000027598 | itch | 1 |
| ENSMUSG00000016024 | Lbp | 0 |
| ENSMUSG00000024052 | Lpin2 | 1 |

| Id | gene_name | Pyroptosis activator 0/ inhibitor 1 |
|---|---|---|
| ENSMUSG00000025779 | Ly96 | 0 |
| ENSMUSG00000032688 | Malt1 | 0 |
| ENSMUSG00000071369 | map3k5 | 0 |
| ENSMUSG00000028284 | map3k7 | 0 |
| ENSMUSG00000053436 | Mapk14 | 0 |
| ENSMUSG00000063065 | mapk3 | 0 |
| ENSMUSG00000016528 | Mapkapk2 | 0 |
| ENSMUSG00000026977 | March7 | 1 |
| ENSMUSG00000037523 | Mavs | 0 |
| ENSMUSG00000022534 | Mefv | 0 |
| ENSMUSG00000019982 | Myb | 0 |
| ENSMUSG00000032508 | Myd88 | 0 |
| ENSMUSG00000078945 | Naip2 | 0 |
| ENSMUSG00000071203 | Naip5 | 0 |
| ENSMUSG00000078942 | Naip6 | 0 |
| ENSMUSG00000028163 | Nfkb1 | 0 |
| ENSMUSG00000025225 | Nfkb2 | 0 |
| ENSMUSG00000021025 | Nfkbia | 1 |
| ENSMUSG00000030595 | Nfkbib | 0 |
| ENSMUSG00000036931 | Nfkbid | 0 |
| ENSMUSG00000023947 | Nfkbie | 0 |
| ENSMUSG00000035356 | Nfkbiz | 0 |
| ENSMUSG00000049871 | Nlrc3 | 1 |
| ENSMUSG00000039193 | Nlrc4 | 0 |
| ENSMUSG00000049709 | Nlrp10 | 1 |
| ENSMUSG00000078817 | Nlrp12 | 1 |
| ENSMUSG00000070390 | Nlrp1b | 0 |
| ENSMUSG00000032691 | Nlrp3 | 0 |
| ENSMUSG00000060508 | Nlrp9b | 0 |
| ENSMUSG00000038058 | Nod1 | 0 |
| ENSMUSG00000055994 | Nod2 | 0 |
| ENSMUSG00000026672 | Optn | 1 |
| ENSMUSG00000038495 | Otud7b | 1 |
| ENSMUSG00000029468 | P2rx7 | 0 |
| ENSMUSG00000030774 | Pak1 | 0 |
| ENSMUSG00000020134 | Peli1 | 0 |
| ENSMUSG00000034462 | Pkd2 | 0 |
| ENSMUSG00000004591 | Pkn2 | 1 |
| ENSMUSG00000020349 | ppp2ca | 1 |
| ENSMUSG00000027843 | Ptpn22 | 0 |
| ENSMUSG00000030793 | Pycard | 0 |
| ENSMUSG00000001847 | Rac1 | 0 |
| ENSMUSG00000027466 | rbck1 | 0 |
| ENSMUSG00000020275 | Rel | 0 |
| ENSMUSG00000024927 | rela | 0 |
| ENSMUSG00000002983 | Relb | 0 |
| ENSMUSG00000021408 | ripk1 | 0 |
| ENSMUSG00000041135 | ripk2 | 0 |
| ENSMUSG00000022221 | Ripk3 | 0 |
| ENSMUSG00000028557 | rnf11 | 0 |

| Id | gene_name | Pyroptosis activator 0/ inhibitor 1 |
|---|---|---|
| ENSMUSG00000047098 | rnf31 | 0 |
| ENSMUSG00000024952 | rps6ka4 | 0 |
| ENSMUSG00000021180 | Rps6ka5 | 0 |
| ENSMUSG00000021180 | Rps6ka6 | 0 |
| ENSMUSG00000050132 | Sarm1 | 1 |
| ENSMUSG00000034252 | Senp6 | 0 |
| ENSMUSG00000022552 | Sharpin | 0 |
| ENSMUSG00000020063 | Sirt1 | 0 |
| ENSMUSG00000015149 | Sirt2 | 0 |
| ENSMUSG00000038037 | socs1 | 1 |
| ENSMUSG00000026104 | Stat1 | 0 |
| ENSMUSG00000040033 | Stat2 | 0 |
| ENSMUSG00000021457 | Syk | 0 |
| ENSMUSG00000015755 | Tab2 | 0 |
| ENSMUSG00000035476 | Tab3 | 0 |
| ENSMUSG00000064289 | Tank | 0 |
| ENSMUSG00000004535 | Tax1bp1 | 1 |
| ENSMUSG00000053477 | Tcf4 | 0 |
| ENSMUSG00000047123 | Ticam1 | 0 |
| ENSMUSG00000056130 | Ticam2 | 0 |
| ENSMUSG00000046688 | Tifa | 0 |
| ENSMUSG00000049625 | Tifab | 1 |
| ENSMUSG00000032041 | tirap | 0 |
| ENSMUSG00000044827 | Tlr1 | 0 |
| ENSMUSG00000051969 | Tlr11 | 0 |
| ENSMUSG00000062545 | Tlr12 | 0 |
| ENSMUSG00000033777 | Tlr13 | 0 |
| ENSMUSG00000027995 | Tlr2 | 0 |
| ENSMUSG00000031639 | Tlr3 | 0 |
| ENSMUSG00000039005 | Tlr4 | 0 |
| ENSMUSG00000051498 | Tlr6 | 0 |
| ENSMUSG00000044583 | Tlr7 | 0 |
| ENSMUSG00000040522 | Tlr8 | 0 |
| ENSMUSG00000024401 | Tnf | 0 |
| ENSMUSG00000019850 | Tnfaip3 | 1 |
| ENSMUSG00000030341 | Tnfrsf1a | 0 |
| ENSMUSG00000028599 | Tnfrsf1b | 0 |
| ENSMUSG00000020400 | Tnip1 | 1 |
| ENSMUSG00000025139 | Tollip | 1 |
| ENSMUSG00000031887 | Tradd | 0 |
| ENSMUSG00000026942 | traf2 | 0 |
| ENSMUSG00000021277 | Traf3 | 1 |
| ENSMUSG00000027164 | Traf6 | 0 |
| ENSMUSG00000021326 | Trim27 | 1 |
| ENSMUSG00000030921 | Trim30a | 1 |
| ENSMUSG00000059552 | Trp53 | 0 |
| ENSMUSG00000009292 | Trpm2 | 0 |
| ENSMUSG00000038393 | Txnip | 0 |
| ENSMUSG00000033685 | ucp2 | 0 |
| ENSMUSG00000031826 | usp10 | 1 |

| Id | gene_name | Pyroptosis activator 0/ inhibitor 1 |
|---|---|---|
| ENSMUSG00000053483 | Usp21 | 1 |
| ENSMUSG00000022710 | usp7 | 0 |
| ENSMUSG00000025860 | xiap | 1 |
| ENSMUSG00000027514 | Zbp1 | 0 |
| ENSMUSG00000044786 | Zfp36 | 0 |
| ENSMUSG00000027540 | Ptpn1 | 1 |
| ENSMUSG00000025199 | Chuk | 0 |

| Id | gene_name | Apoptosis activator 0/ inhibitor 1 |
|---|---|---|
| ENSMUSG00000026842 | Abl1 | 1 |
| ENSMUSG00000022185 | acin1 | 0 |
| ENSMUSG00000018500 | Adora2b | 0 |
| ENSMUSG00000036932 | aifm1 | 0 |
| ENSMUSG00000024847 | AIP | 1 |
| ENSMUSG00000001729 | akt1 | 1 |
| ENSMUSG00000004056 | akt2 | 1 |
| ENSMUSG00000019699 | Akt3 | 1 |
| ENSMUSG00000019979 | Apaf1 | 0 |
| ENSMUSG00000027193 | Api5 | 1 |
| ENSMUSG00000010911 | apip | 1 |
| ENSMUSG00000022602 | Arc | 1 |
| ENSMUSG00000026628 | Atf3 | 0 |
| ENSMUSG00000042406 | Atf4 | 0 |
| ENSMUSG00000032905 | Atg12 | 1 |
| ENSMUSG00000022663 | Atg3 | 1 |
| ENSMUSG00000020897 | Aurkb | 1 |
| ENSMUSG00000003604 | Aven | 1 |
| ENSMUSG00000024959 | bad | 0 |
| ENSMUSG00000037316 | Bag4 | 1 |
| ENSMUSG00000057789 | Bak1 | 0 |
| ENSMUSG00000003873 | Bax | 0 |
| ENSMUSG00000002083 | Bbc3 | 0 |
| ENSMUSG00000028191 | Bcl10 | 0 |
| ENSMUSG00000057329 | Bcl2 | 1 |
| ENSMUSG00000089929 | Bcl2a1b | 1 |
| ENSMUSG00000053820 | Bcl2a1c | 1 |
| ENSMUSG00000007659 | bcl2l1 | 1 |
| ENSMUSG00000027381 | Bcl2l11 | 0 |
| ENSMUSG00000003190 | Bcl2l12 | 1 |
| ENSMUSG00000009112 | Bcl2l13 | 1 |
| ENSMUSG00000089682 | bcl2l2 | 1 |
| ENSMUSG00000004446 | Bid | 0 |
| ENSMUSG00000016758 | bik | 0 |
| ENSMUSG00000057367 | Birc2 | 1 |
| ENSMUSG00000032000 | birc3 | 1 |
| ENSMUSG00000017716 | Birc5 | 1 |
| ENSMUSG00000024073 | Birc6 | 1 |
| ENSMUSG00000021936 | bmf | 0 |
| ENSMUSG00000049086 | Bmyc | 1 |
| ENSMUSG00000078566 | Bnip3 | 0 |
| ENSMUSG00000022051 | Bnip3l | 0 |
| ENSMUSG00000026278 | bok | 0 |
| ENSMUSG00000002413 | Braf | 1 |
| ENSMUSG00000020423 | Btg2 | 0 |
| ENSMUSG00000024942 | Capn1 | |
| ENSMUSG00000026509 | Capn2 | 0 |
| ENSMUSG00000025887 | Casp12 | 0 |
| ENSMUSG00000029863 | Casp2 | 0 |

| Id | gene_name | Apoptosis activator 0/ inhibitor |
|---|---|---|
| ENSMUSG00000031628 | Casp3 | 0 |
| ENSMUSG00000027997 | Casp6 | 0 |
| ENSMUSG00000025076 | Casp7 | 0 |
| ENSMUSG00000026029 | Casp8 | 0 |
| ENSMUSG00000028914 | casp9 | 0 |
| ENSMUSG00000021585 | Cast | 1 |
| ENSMUSG00000035042 | ccl5 | 1 |
| ENSMUSG00000079227 | Ccr5 | 1 |
| ENSMUSG00000025358 | cdk2 | 0 |
| ENSMUSG00000044303 | cdkn2a | 0 |
| ENSMUSG00000026031 | Cflar | 1 |
| ENSMUSG00000032113 | chek1 | 1 |
| ENSMUSG00000024526 | cidea | 0 |
| ENSMUSG00000022219 | cideb | 0 |
| ENSMUSG00000040782 | cop1 | 1 |
| ENSMUSG00000045867 | CRADD | 0 |
| ENSMUSG00000014599 | Csf1 | 1 |
| ENSMUSG00000024621 | Csf1r | 1 |
| ENSMUSG00000006932 | Ctnnb1 | 1 |
| ENSMUSG00000021939 | Ctsb | 0 |
| ENSMUSG00000063694 | cycs | 0 |
| ENSMUSG00000002307 | Daxx | 0 |
| ENSMUSG00000030641 | Ddias | 1 |
| ENSMUSG00000025408 | Ddit3 | 0 |
| ENSMUSG00000028974 | Dffa | 0 |
| ENSMUSG00000029433 | diablo | 0 |
| ENSMUSG00000014773 | Dll1 | 1 |
| ENSMUSG00000022789 | Dnm1l | |
| ENSMUSG00000024137 | e4f1 | 1 |
| ENSMUSG00000015337 | endog | 0 |
| ENSMUSG00000031077 | fadd | 0 |
| ENSMUSG00000024778 | Fas | 0 |
| ENSMUSG00000000817 | faslg | 0 |
| ENSMUSG00000035764 | fbxo45 | 1 |
| ENSMUSG00000021250 | Fos | 0 |
| ENSMUSG00000003545 | Fosb | 0 |
| ENSMUSG00000024912 | Fosl1 | 0 |
| ENSMUSG00000029135 | fosl2 | 0 |
| ENSMUSG00000048756 | foxo3 | 0 |
| ENSMUSG00000042903 | Foxo4 | 0 |
| ENSMUSG00000007989 | fzd3 | 1 |
| ENSMUSG00000036390 | Gadd45a | 1 |
| ENSMUSG00000015312 | Gadd45b | 1 |
| ENSMUSG00000021453 | Gadd45g | 1 |
| ENSMUSG00000030498 | gas2 | 1 |
| ENSMUSG00000031451 | Gas6 | 1 |
| ENSMUSG00000051136 | ghsr | 1 |
| ENSMUSG00000022812 | gsk3b | 0 |
| ENSMUSG00000028800 | Hdac1 | 1 |
| ENSMUSG00000061436 | Hipk2 | 0 |

| Id | gene_name | Apoptosis activator 0/ inhibitor 1 |
|---|---|---|
| ENSMUSG00000066551 | Hmgb1 | 0 |
| ENSMUSG00000046607 | hrk | 0 |
| ENSMUSG00000020048 | Hsp90b1 | 1 |
| ENSMUSG00000004951 | Hspb1 | 1 |
| ENSMUSG00000068329 | htra2 | 0 |
| ENSMUSG00000020053 | Igf1 | 1 |
| ENSMUSG00000005533 | Igf1r | 1 |
| ENSMUSG00000027598 | Itch | 1 |
| ENSMUSG00000052684 | jun | 0 |
| ENSMUSG00000052837 | Junb | 0 |
| ENSMUSG00000071076 | jund | 0 |
| ENSMUSG00000000708 | kat2b | 1 |
| ENSMUSG00000024926 | kat5 | 1 |
| ENSMUSG00000036940 | Kdm1a | 1 |
| ENSMUSG00000049327 | kmt5a | 1 |
| ENSMUSG00000030265 | Kras | 1 |
| ENSMUSG00000034394 | Lif | 0 |
| ENSMUSG00000028063 | lmna | 1 |
| ENSMUSG00000033352 | Map2k4 | 1 |
| ENSMUSG00000002948 | Map2k7 | 1 |
| ENSMUSG00000020941 | Map3k14 | 0 |
| ENSMUSG00000014426 | Map3k4 | 1 |
| ENSMUSG00000071369 | Map3k5 | 0 |
| ENSMUSG00000053436 | Mapk14 | 1 |
| ENSMUSG00000021936 | mapk8 | 0 |
| ENSMUSG00000020366 | Mapk9 | 0 |
| ENSMUSG00000038612 | Mcl1 | 1 |
| ENSMUSG00000020184 | Mdm2 | 1 |
| ENSMUSG00000054387 | Mdm4 | 1 |
| ENSMUSG00000005583 | Mef2c | 0 |
| ENSMUSG00000027668 | Mfn1 | 1 |
| ENSMUSG00000029020 | Mfn2 | 0 |
| ENSMUSG00000032591 | Mst1 | 0 |
| ENSMUSG00000025025 | Mxi1 | 1 |
| ENSMUSG00000022346 | Myc | 0 |
| ENSMUSG00000026393 | nek7 | 1 |
| ENSMUSG00000000120 | Ngfr | 1 |
| ENSMUSG00000055994 | Nod2 | 1 |
| ENSMUSG00000027852 | Nras | 1 |
| ENSMUSG00000024975 | Pdcd4 | 0 |
| ENSMUSG00000006494 | Pdk1 | 1 |
| ENSMUSG00000055866 | Per2 | 0 |
| ENSMUSG00000032405 | Pias1 | 0 |
| ENSMUSG00000025507 | Pidd1 | 0 |
| ENSMUSG00000027665 | Pik3ca | 1 |
| ENSMUSG00000032462 | Pik3cb | 1 |
| ENSMUSG00000039936 | Pik3cd | 1 |
| ENSMUSG00000041417 | Pik3r1 | 1 |
| ENSMUSG00000031834 | Pik3r2 | 1 |
| ENSMUSG00000028698 | Pik3r3 | 1 |

| Id | gene_name | Apoptosis activator 0/ inhibitor 1 |
|---|---|---|
| ENSMUSG00000024014 | Pim1 | 1 |
| ENSMUSG00000032171 | pin1 | 0 |
| ENSMUSG00000050721 | Plekho2 | 0 |
| ENSMUSG00000030867 | Plk1 | 1 |
| ENSMUSG00000021701 | Plk2 | 1 |
| ENSMUSG00000024521 | Pmaip1 | 0 |
| ENSMUSG00000020525 | ppm1d | 1 |
| ENSMUSG00000021285 | Ppp1r13b | 0 |
| ENSMUSG00000007564 | Ppp2r1a | 0 |
| ENSMUSG00000032058 | Ppp2r1b | 0 |
| ENSMUSG00000022092 | Ppp3cc | 0 |
| ENSMUSG00000050697 | Prkaa1 | 0 |
| ENSMUSG00000029513 | Prkab1 | 0 |
| ENSMUSG00000038205 | Prkab2 | 1 |
| ENSMUSG00000023110 | prmt5 | 1 |
| ENSMUSG00000013663 | pten | 0 |
| ENSMUSG00000001847 | Rac1 | 0 |
| ENSMUSG00000010067 | Rassf1 | 0 |
| ENSMUSG00000029397 | Rchy1 | |
| ENSMUSG00000038555 | Reep2 | |
| ENSMUSG00000020696 | Rffl | 1 |
| ENSMUSG00000021408 | Ripk1 | 0 |
| ENSMUSG00000029474 | Rnf34 | 1 |
| ENSMUSG00000024290 | rock1 | 0 |
| ENSMUSG00000023809 | Rps6ka2 | 1 |
| ENSMUSG00000075701 | selenos | 1 |
| ENSMUSG00000037111 | setd7 | 0 |
| ENSMUSG00000042557 | Sin3a | 0 |
| ENSMUSG00000020063 | Sirt1 | 1 |
| ENSMUSG00000064326 | Siva1 | 0 |
| ENSMUSG00000016319 | Slc25a5 | 0 |
| ENSMUSG00000028645 | Slc2a1 | 1 |
| ENSMUSG00000003153 | Slc2a3 | 1 |
| ENSMUSG00000032402 | Smad3 | 0 |
| ENSMUSG00000026603 | Smyd2 | 1 |
| ENSMUSG00000001280 | sp1 | 0 |
| ENSMUSG00000022885 | st6gal1 | 1 |
| ENSMUSG00000030224 | Strap | 1 |
| ENSMUSG00000039615 | stub1 | 0 |
| ENSMUSG00000024807 | syvn1 | 1 |
| ENSMUSG00000004535 | Tax1bp1 | 1 |
| ENSMUSG00000068039 | tcp1 | 0 |
| ENSMUSG00000028958 | tmub1 | 0 |
| ENSMUSG00000024401 | Tnf | 0 |
| ENSMUSG00000019850 | Tnfaip3 | 1 |
| ENSMUSG00000022074 | Tnfrsf10b | 0 |
| ENSMUSG00000030341 | Tnfrsf1a | 0 |
| ENSMUSG00000028599 | Tnfrsf1b | 0 |
| ENSMUSG00000039304 | Tnfsf10 | 0 |
| ENSMUSG00000036822 | topors | 1 |
| ENSMUSG00000031887 | Tradd | 0 |
| ENSMUSG00000026942 | traf2 | 0 |
| ENSMUSG00000026637 | Traf5 | 0 |
| ENSMUSG00000029833 | Trim24 | 1 |
| ENSMUSG00000059552 | Trp53 | 0 |
| ENSMUSG00000026510 | trp53bp2 | 0 |
| ENSMUSG00000035186 | Ubd | 1 |
| ENSMUSG00000031826 | usp10 | 1 |
| ENSMUSG00000022710 | usp7 | 1 |
| ENSMUSG00000041058 | Wwp1 | 1 |
| ENSMUSG00000053110 | Yap1 | 1 |
| ENSMUSG00000027663 | Zmat3 | 0 |
| ENSMUSG00000021640 | Naip1 | 1 |
