## Supplementary Table 2 for "*Leishmania amazonensis* controls macrophage-regulated cell death to establish chronic infection *in vitro* and *in vivo*"

**Lecoeur et al., Supplementary Table 2**

| | Forward | Reverse |
|---|---|---|
| **Akt3** | TGATGGAATATGTTAATGGCGGA | GAGATCACGGTACACAATCTT |
| **Bax** | ATATGGAGCTGCAGAGGA | CAAAGTAGAAGAGGGCAACC |
| **Bcl2l11** | GACAGTCTCAGGAGGAACC | GTGTAAGTTTCGTTGAACTCG |
| **Birc2** | ATGCTGACCCTACAGAGAC | CACCAGGCTCCTACTGAA |
| **Birc3** | GGTGGAACATGCCAAGTG | GTCTGACGTAGATAATAGCTGC |
| **Birc6** | GTGGTTCAGCTGGAACT | TTAAACCATCTTTGAGCTGTGT |
| **Birc6** | GTGGTTCAGCTGGAACT | TTAAACCATCTTTGAGCTGTGT |
| **Camk2d** | TCACGACAGTATATCGGAGGA | ATACAATGACTGGCATCAGC |
| **Casp1** | ATCTGTATTCACGCCCT | ATCTGTATTCACGCCCT |
| **Casp2** | GCCTATCCACAGATGCTACG | TCAGGAGTGCATGGCTT |
| **Casp3** | ATTAATGGATAGTGTTTCTAAGGAAGAT | AGTCAGACTCCGGCAGTA |
| **Casp8** | AAATGTAAGCTGGAAGATGACT | CCAGCAGGCTCTTGTTG |
| **Cd14** | GCCAAATTGGTCGAACA | CTCGTCTAGCTCGCAG |
| **Cflar** | TGGAGAAACTGAATCTAATTGCTT | GATCTTGCTCCTTGGCTG |
| **Ddx58** | GAGGATGATGGAGCGGA | GGTTTCAATGGGCTGTGTAA |
| **Fadd** | GCCTGGACGACTTCGAG | GCCTGGACGACTTCGAG |
| **Fbxl2** | GTGCGGTGGCTTCCTTA | GTGCGGTGGCTTCCTTA |
| **Ifnar1** | TAAAGTGGAGCAGCCACG | TAAAGAGAATTCACACTTGGTCG |
| **Il18** | GCTGGAGACCTGGAAT | GCTGGAGACCTGGAAT |
| **Itch** | GTCACAGTAGATGGACAGTCAA | GTCACAGTAGATGGACAGTCAA |
| **Map3k5** | ACCTGTTGCTTTCCTACAGA | GTTCAGTGCAAATGCGTAAT |
| **Mcl1** | AAACACTTAAAGAGCGTAAACCAA | CCAGCAGCACATTTCTGA |
| **Mefv** | CTGGATGAGATGATTGAAGAACTAGA | AGTGTCCAACAGCTCAG |
| **Mlkl** | GCAGCAGGAAGATCGAC | TGTGGGATCTCCTGTGT |
| **Nfkb1** | GCAACTCACAGACAGAGAGAA | CTGTGAACATGAGGCGCA |
| **Nfkb2** | GCTTCAGATTTCGATATGGCT | GCCGGTCCCTCATAGTT |
| **Nlrc3** | GCTGGACAACAACCAGTTC | GGGCTTTGGCTCCTTTGTTA |
| **Nlrc4** | GTCTCACACTCAAAGGG | GTCTCACACTCAAAGGG |
| **Nlrp3** | GACTTTGGAATCAGATTGCT | GGGTCCTTCATCTTTTCAC |
| **Nod1** | CCGTCTCACGGTTATCAG | CCGTCTCACGGTTATCAG |
| **Optn** | CTTCGTTGAGATCAGGATGAC | CTTCGTTGAGATCAGGATGAC |
| **P2rx7** | TGAGCGATAAGCTGTACC | TGAGCGATAAGCTGTACC |
| **Pdk1** | TCATCGAAAGCACATTGGAAG | TTCAAGTTCAGGAGAGTTAACATAATAC |
| **Pidd** | TATCCCAGGTTCCAGAAATGC | GCTCCTGCTGGAAACTGTA |

| | | |
|---|---|---|
| **Pik3cb** | CTAATGTGTCAAGTCGTGGT | CTTGCCGTAGAGTCCAAATA |
| **Pim** | CTGTCCAAGATCAACTCCC | GGCTCCTTCTCTTTGCC |
| **Pycard** | TGCTTAGAGACATGGGC | TGCTTAGAGACATGGGC |
| **Rela** | CCTTTCAATGGACCAACTG | TGATGGTGCTGAGGGAT |
| **Rffl** | GGTGACTCGGCTGTACAA | GGTGAGTCCATGCAAATCT |
| **Ripk1** | CCCAGATAGATGTCCCACT | CCCAGATAGATGTCCCACT |
| **Ripk3** | GGGTTCCCAGTCCGAAA | CTTCTGTGCTGAGACAGATAAT |
| **Tax1bp1** | CAACAGTCAATTGTGTACTAGC | GAACTGGAAAGGAGTACTTGC |
| **Tlr4** | TCTTCTCCTGCCTGACAC | TGGTTGAAGAAGGAATGTCATC |
| **Tlr8** | ACCAGTGCCATCTTCCATA | TTCTGCAATCACAAGGGAGT |
| **Tnfaip3** | CTGAAATCTCAGGAATTTGTGG | CAGGTGTGTCTGCTGATG |
| **Tnfr1a** | TCCGCTTGCAAATGTCAC | GGGATATCGGCACATTAAACT |
| **Tnip1** | CTGCGCCAGGAGAATGA | CTGCGCCAGGAGAATGA |
| **Tradd** | GAGCAGCTGGACAGTTG | GAGCAGCTGGACAGTTG |
| **Trp53** | CTCTGAGCCAGGAGACA | CTTCACTTGGGCCTTCAAA |
| **Usp2** | TTCCTTCGTTTCCTTCTGGATG | TCAAGGGTCTCAGGGCT |
| **Zbp1** | GCTCATTCTGGAGTCACAC | TTGGCTCCTTGTTGGCA |
